# The lncRNA *Malat1* is trafficked to the cytoplasm as a localized mRNA encoding a small peptide in neurons

**DOI:** 10.1101/2024.02.01.578240

**Authors:** Wen Xiao, Reem Halabi, Chia-Ho Lin, Mohammad Nazim, Kyu-Hyeon Yeom, Douglas L Black

**Author notes:** **Author Contributions:** W.X., and D.L.B. designed the research; W.X., R.H., K-H.Y., and M.N. performed the experiments; W.X. and C.-H. L. analyzed the data; and W.X. and D.L.B. wrote the paper. **Competing Interest Statement:** D.L.B. has equity and serves on the board of directors for Panorama Medicine. This company did not contribute to or direct any of the research reported in this article.

## Abstract

Synaptic function is modulated by local translation of mRNAs that are transported to distal portions of axons and dendrites. The Metastasis-associated lung adenocarcinoma transcript 1 (*MALAT1*) is broadly expressed across cell types, almost exclusively as a nuclear non-coding RNA. We found that in differentiating neurons, a portion of *Malat1* RNA redistributes to the cytoplasm. Depletion of *Malat1* from neurons stimulated expression of particular pre- and post-synaptic proteins, implicating *Malat1* in their regulation. Neuronal *Malat1* is localized to both axons and dendrites in puncta that co-stain with Staufen1 protein, similar to neuronal granules formed by locally translated mRNAs. Ribosome profiling of mouse cortical neurons identified ribosome footprints within a region of *Malat1* containing short open reading frames. The upstream-most reading frame (M1) of the *Malat1* locus was linked to the GFP coding sequence in mouse ES cells. When these gene-edited cells were differentiated into glutamatergic neurons, the M1-GFP fusion protein was expressed. Antibody staining for the M1 peptide confirmed its presence in wildtype neurons, and showed enhancement of M1 expression after synaptic stimulation with KCL. Our results indicate that *Malat1* serves as a cytoplasmic coding RNA in the brain that is both modulated by and modulates synaptic function.

## Introduction

Long non-coding RNAs (lncRNAs) are RNA molecules longer than ∼500 nucleotides (nt) that lack extended open reading frames (Mattick et al. 2023; Ransohoff et al. 2018). LncRNAs localized to the nucleus can function in chromatin organization, nuclear architecture, genome stability, transcriptional regulation, and RNA processing (Böhmdorfer and Wierzbicki 2015; Khanduja et al. 2016; Tang et al. 2017; Bergmann and Spector 2014; Ouyang et al. 2022), while cytoplasmic lncRNAs play similarly diverse roles in RNA stability, microRNA and protein sequestration, and translational control (Noh et al. 2018; Munschauer et al. 2018; Lee et al. 2016; Karakas and Ozpolat 2021). Despite their noncoding classification, many cytoplasmic lncRNAs have been found to associate with ribosomes and be translated (Ingolia et al. 2014; Ruiz-Orera et al. 2014; Wang et al. 2016; Xing et al. 2021). Short peptides encoded by lncRNA open reading frames (micro ORFs) were found to have function in mRNA processing, DNA repair, muscle regeneration and development, and cancer progression (Anderson et al. 2015; Nelson et al. 2016; Zhang et al. 2017; Bi et al. 2017; Matsumoto et al. 2017; Huang et al. 2017; Zhang et al. 2022). Many new micropeptides were recently identified in human brain, although their roles in neuronal maturation or activity are mostly unknown (Duffy et al. 2022).

*Malat1* (Metastasis Lung cancer Associated Transcript 1) is an abundant and highly conserved lncRNA expressed in many mammalian cell types. The major *Malat1* transcript (∼7-kb in humans and 6.7-kb in mouse) lacks introns and a poly (A) tail, unlike a typical mRNA. Instead the Malat1 transcript undergoes an unusual 3’ end processing reaction where it is cleaved by RNase P to generate a tRNA-like small RNA (mascRNA), which is transported to the cytoplasm (Wilusz et al. 2008, 2012; Brown et al. 2012). The 5’ major portion of the cleaved transcript forms a triple helical structure at its 3’ end that protects it from degradation. These mature *Malat1* transcripts are enriched in nuclear speckles, and have been found to affect splicing, chromatin organization and transcription (Tripathi et al. 2010; Engreitz et al. 2014; Chen et al. 2017; Miao et al. 2022). In neurons, depletion of *Malat1* was found to reduce expression of synaptic proteins and to reduce neurite outgrowth. These effects were attributed to changes in transcription or miRNA availability mediated by the nuclear Malat1 RNA (Bernard et al. 2010; Chen et al. 2016; Kim et al. 2018; Xie et al. 2021). However, a recent study identified m6A modified Malat1 RNA at neuronal synapses and reported that Malat1 depletion impaired fear extinction memory.

In this study, we report that *Malat1* transcripts are exported to the cytoplasm and transported into neuronal processes during neuronal development. Unlike previous observations, we found that depletion of *Malat1* from neurons led to upregulation of pre- and post-synaptic proteins important for neuronal maturation. We demonstrate that *Malat1* colocalizes with the neuronal granule protein Staufen1 in puncta within both axons and dendrites of mature neurons. We further discovered that this neuronal MALAT1 is translated to produce a micropeptide, and that expression of this micropeptide is stimulated by synaptic activity. These findings suggest alternative mechanisms for how the *Malat1* RNA can affect neuronal maturation and activity.

## Results

### A portion of *Malat1* RNA is exported to the cytoplasm in differentiating neurons and transported into neurites

*Malat1* is a well-studied lncRNA enriched in nuclear speckles and largely absent from the cytoplasm across many cell types (Tripathi et al. 2010; Nakagawa et al. 2012; Miyagawa et al. 2012). We previously generated extensive RNA-sequencing data from fractionated cellular compartments (Yeom et al. 2021). Total rRNA-depleted RNA was extracted and sequenced from chromatin, nucleoplasm and cytoplasm fractions of three mouse cell types: ES cells (mESC), neuronal progenitor cells (NPC), and primary cortical neurons explanted from 15 day embryos and differentiated for 5 days in culture. We observed that, in contrast to ESC and NPC, cortical neurons displayed abundant *Malat1* RNA in the cytoplasm in addition to that in the nuclear fractions (Fig. 1A). Other nuclear lncRNAs, including *Neat1* (Yeom et al. 2021) and *kcnq1ot1* (Supplemental Fig. 1A) maintained their almost exclusively nuclear expression in all three cell types.

**Fig. 1:**
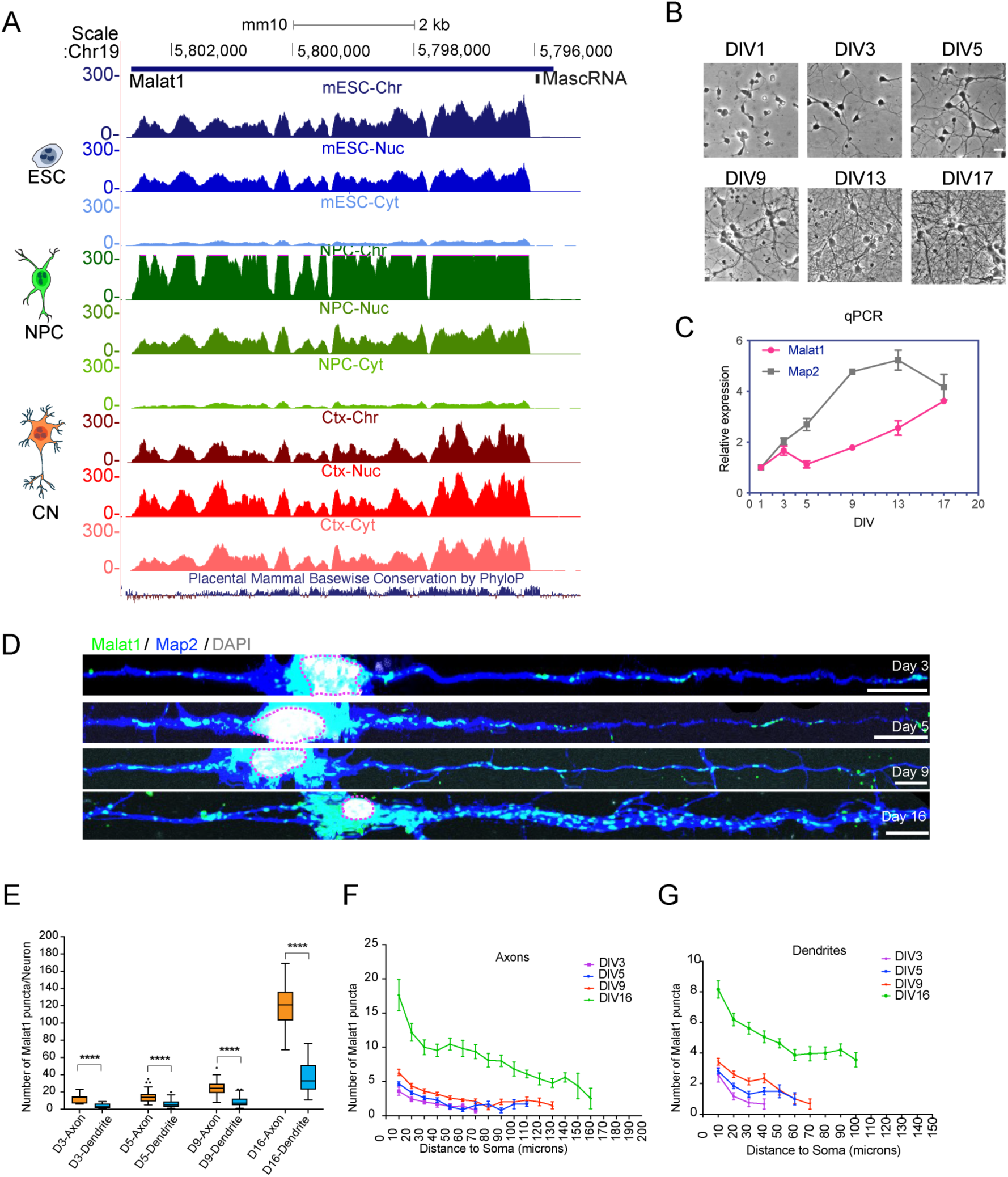
*Malat1* is exported from the nucleus to the cytoplasm during neuronal differentiation. (A) Genome browser tracks of the *Malat1* locus displaying RNAseq reads from chromatin, nucleoplasmic, and cytoplasmic fractions of three cell types: Blue, mouse embryonic stem cells; Green, neuronal progenitor cells; Red, primary cortical neurons at DIV5. (B) Morphology of cultured primary cortical neurons at different DIVs used for RNA quantification in Fig. 1C. (C) RT/qPCR analysis of *Malat1* and *Map2* expression in the cells shown in Fig. 1B. (D) *Malat1* RNA FISH (green) combined with Map2 protein staining (blue) in neurons at different stages of development. Scale bar: 10 um. (E) Quantification of *Malat1* FISH puncta in axons and neurites. (F) and (G), *Malat1* spot counts along axons and dendrites with distance from cell body (soma) at different DIVs. “****” indicates P value < 0.0001.

We cultured primary cortical neurons from E16 mice (Fig 1B) and quantified *Malat1* RNA abundance across neuronal maturation by RT/qPCR. We found *Malat1* expression increased 4-fold relative to *Gapdh* between DIV0 and DIV17 (Fig. 1C). This roughly paralleled a 5-fold increase in the neuronal mRNA *Map2* although the two transcripts differed in their abundance profiles across time (Fig. 1C).

Cell nuclei are difficult to cleanly isolate from cultured neurons after about 5 days of in vitro culture (DIV5). To confirm the release of *Malat1* into the cytoplasm of mature neurons, we used a digitonin elution assay at various stages of neuronal maturation (Supplemental Fig. 1B) to permeabilize the plasma membrane and selectively release the cytoplasmic components (Adam 2016; Niklas et al. 2011). This material was compared to the remaining cellular material containing both cytoplasmic and nuclear contents. RNA was then extracted from the two fractions and assayed by reverse transcription PCR (RT-PCR) (Supplemental Fig. 1B). Notably, cytoplasmic *Malat1* transcripts were detected at DIV 2 and increased at DIV 5 and 10 (Supplemental Fig. 1B). This was similar to cytoplasmic mRNAs (*Gapdh* and *Actb*) and in contrast to the nuclear RNAs *Neat1* and *U6* that were found almost entirely in the combined nuclear and cytoplasmic fraction at all DIVs (Supplemental Fig. 1B).

We also analyzed previously published RNAseq data generated from rat neuronal processes that had extended through a filter to allow the clean separation of cell projections from cell bodies and nuclei (Saini et al. 2019). These data showed that *Malat1* was abundantly expressed in neurites, whereas another nuclear lncRNA *Neat1* was absent (Supplemental Fig. 1C, D)(Saini et al. 2019).

We next sought to directly observe *Malat1* in neurons and assess its subcellular distribution using single molecule RNA fluorescence in situ hybridization (smFISH). We designed and labeled 94 fluorescently tagged oligonucleotides that tile the *Malat1* sequence (Xiao et al. 2023). As expected, these probes densely stained the nuclei in neurons throughout maturation (Fig. 1D). In addition, there were many small *Malat1* stained puncta in the neuronal processes, whose number increased with maturation (Fig. 1D, Fig. 1E). *Malat1* puncta were observed in both axons and dendrites, with larger numbers in axons, as defined by the cellular morphology (Fig. 1E). Measuring the punctal density of *Malat1* along axons or dendrites, the *Malat1* puncta decreased with distance from the soma (cell body) (Fig. 1F-G). To confirm the specificity of the *Malat1* FISH signal, we split the 94 *Malat1* probes into two subsets targeting either the 5’ or 3’ portion of the *Malat1* transcript. Each 47-probe subset was labeled with a different fluorophore (ATTO565 or ATTO647N) (Supplemental Fig. 2A). As shown in Supplemental Fig. 2B,C the two probe subsets co-localized well along the neurites, indicating they are staining both the 5’ and 3’ portions of the RNA. Overall these results demonstrate that *Malat1* RNA is transported from the nucleus to the cytoplasm in developing cortical neurons and is trafficked into neurites away from the soma.

### Depletion of *Malat1* stimulates the expression of synaptic proteins

Previous studies found that *Malat1* depletion by ASOs decreased the levels of synapsin and other synaptic markers in cultured hippocampal neurons, while *Malat1* overexpression led to increases in synapse density (Bernard et al. 2010; Madugalle et al. 2023). These effects were thought to result from the loss of nuclear *Malat1*, but given the observations above, they could also result from effects of cytoplasmic *Malat1*. To examine the effects of *Malat1* in our system, we designed three GapmeR oligonucleotides (ASO-a, ASO-b and ASO-c) complementary to sequences in *Malat1* that target the RNA for degradation by RNase H (Supplemental Fig. 2D). Each of these ASO’s induced efficient (>90%) depletion of *Malat1* measured by reverse transcription-qPCR (Supplemental Fig.2E), and eliminated *Malat1* staining by FISH (Supplemental Fig. 2F). To examine the effects of *Malat1* depletion in our cortical cultures, we treated the cells with Gapmer ASO’s, and then assayed a variety of neuronal and synaptic markers by RT/qPCR and immunoblot. Surprisingly, we observed increased mRNA levels for some synaptic proteins such as Synaptophysin and NRGN, and for the neuronal beta-tubulin protein TuJ1 (Fig. 2A). The magnitude of these mRNA changes varied depending on the protein, with the largest being about two-fold for NRGN. Synaptophysin and PSD95 proteins also increased about two-fold upon *Malat1* depletion, as measured by immunoblot (Fig. 2B-C). This was confirmed by immunofluorescence, where both the presynaptic synaptophysin and postsynaptic PSD95 were seen to increase in the soma and throughout the dendritic arbor after Gapmer treatment (Fig. 2D-G). Thus in our system, *Malat1* acts to reduce the expression of certain neuronal proteins. Why *Malat1* depletion in our system might have the opposite effect of the previous observations is not clear.

**Fig. 2:**
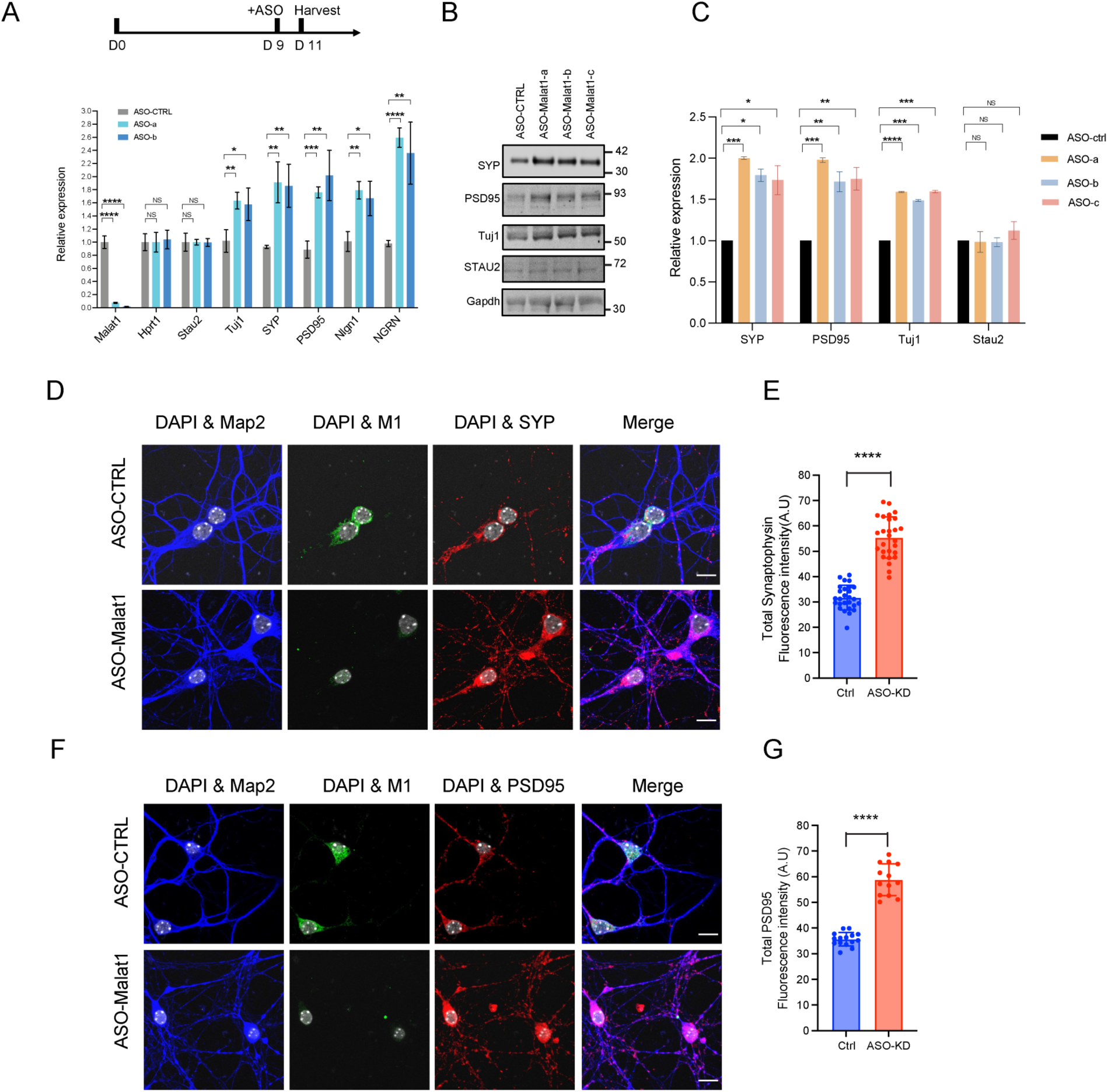
*Malat1* knockdown stimulates expression of pre- and post-synaptic proteins. A) Top, timeline of ASO treatment in neuronal culture. Bottom. qPCR analysis of RNA levels for selected genes after *Malat1* knockdown compared to control (Ctrl) ASO treatment. (B) Immunoblot analysis of *SYP*, *PSD95*, *Tuj1*, *Stau2* and *Gapdh* proteins after *Malat1* knockdown. Immunofluorescent secondary antibodies were employed to detect and quantify each protein signal. (C) Quantification of immunoblot bands in b measured relative to Gapdh. (D) Immunofluorescence of Synaptophysin and M1 peptide before and after *Malat1* depletion by ASOs. (E) Quantification of total Synaptophysin fluorescense intensity in Control (CTRL) and *Malat1* KD neurons. (F) Immunofluorescence of PSD95 and M1 peptide after *Malat1* depletion by ASOs. (G) Quantification of mean PSD95 intensity in Control and *Malat1* KD neurons. “*” indicates a P value ≤ 0.05; “**” P value ≤ 0.01; “***” P value ≤ 0.001; “****” P value < 0.0001, “NS” indicates an nonsignificant P value > 0.05.

### *Malat1* co-localizes with Staufen1 in neuronal mRNA granules

Many neuronal mRNAs are packaged into mRNP granules for their transport into distal processes where they are locally translated (Bauer et al. 2023; Grzejda et al. 2022; Knowles et al. 1996; Holt et al. 2019). The mRNAs in these particles are densely packaged with proteins and have been observed to have low accessibility to FISH probes (Bauer et al. 2023; Fritzsche et al. 2013). Limited proteinase treatment has been employed to expose these RNAs and boost their FISH signals (Young et al. 2020; Sato et al. 2022; Buxbaum et al. 2014). Similar to the previous observations, we found that, *Malat1* puncta in the cytoplasm became brighter and more numerous after limited proteinase K treatment (Fig. 3A-B). In contrast, the FISH signal for the actively translated *Gapdh* mRNA was not altered by the protease treatment (Fig. 3A-B).

**Fig. 3:**
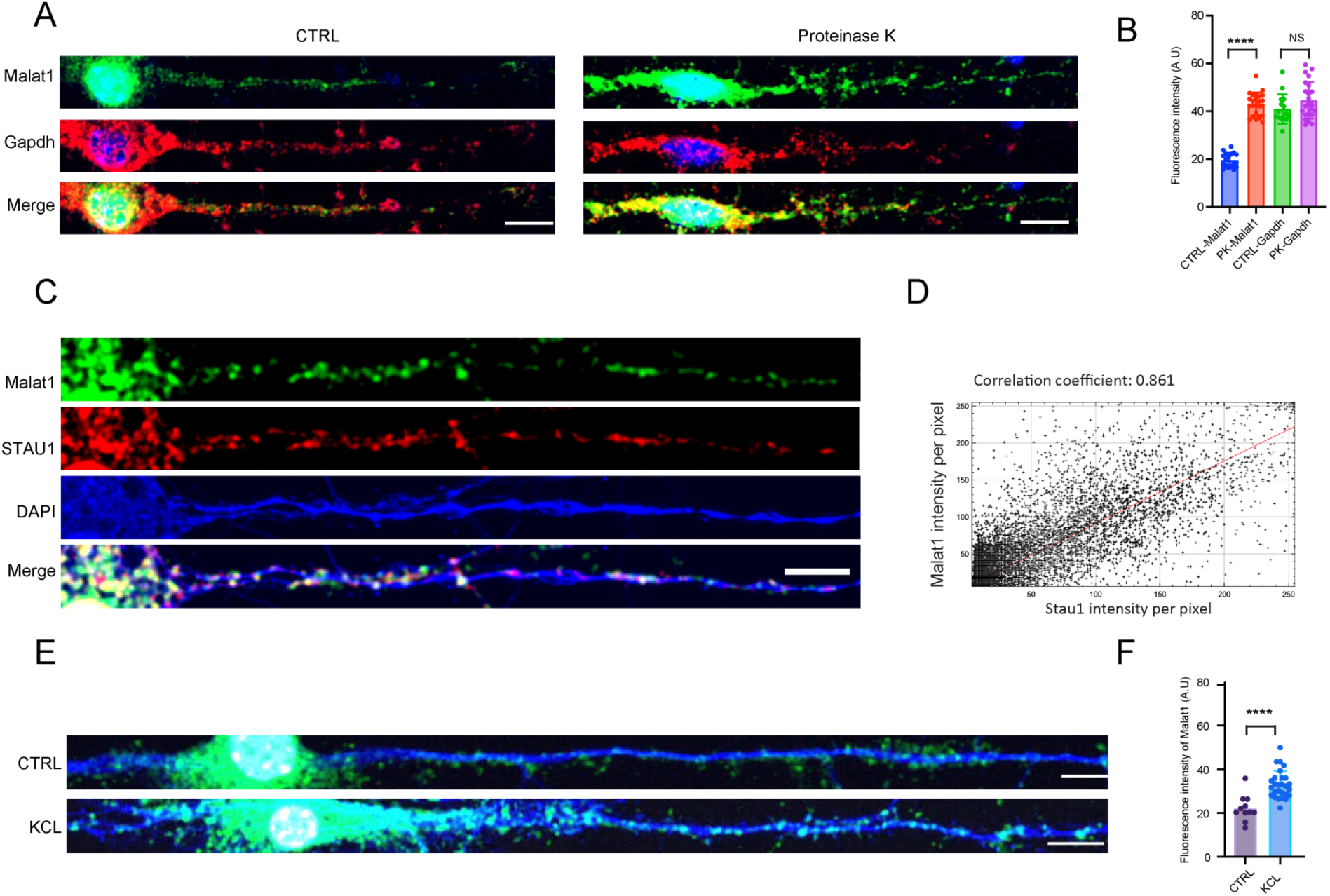
*Malat1* RNA is masked by protein in the cytoplasm and costains with Staufen1. (A) Left, *Malat1* (green) and Gapdh (red) RNA FISH in control primary neurons at DIV 13. Right, *Malat1* (green) and *Gapdh* (red) RNA FISH in neurons treated with limited proteinase K (see methods). (B) Quantification of the mean fluorescent intensity in cells shown in a. 25 cells were measured for each probe and condition. (C) *Malat1* RNA FISH combined with Map2 and STAU1 protein staining in cultured cortical neurons at DIV 13. (D) Pixel intensity correlation of the Stau1 and *Malat1* signals. (E) RNA FISH of *Malat1* (green) in Control primary neurons (H2O) and neurons exposed to 100 mM KCL for 60 minutes, with fixation 10 minutes later. Blue, Map2 protein stain. (F) Mean fluorescent intensity quantification for 20 cells of each condition in e. Scale bar, 10 um.

One protein associated with mRNA in neuronal granules is Staufen1 (Mallardo et al. 2003; Kiebler and Bassell 2006). To examine Staufen1 association with *Malat1*, we combined smFISH and Immunofluorescence (IF) using the *Malat1* hybridization probes and an antibody against Staufen1. We found *Malat1* and Staufen1 were strongly but not completely colocalized in the cytoplasm of neurons at DIV16 (Fig. 3C-D). The staining of the related Staufen2 protein was also strongly correlated with the *Malat1* FISH signal (Supplemental Fig. 3A-B). In contrast, CaMK2a mRNA, a locally translated mRNA that forms granules, showed minimal overlap with *Malat1*, indicating that these two RNAs are in different granules (Supplemental Fig. 3C-D). Neuronal mRNA granules are trafficked along neuronal processes but are excluded from synaptic spines (Kiebler and Bassell 2006; Batish et al. 2012). Costaining for the pre- and postsynaptic proteins Synaptophysin and PSD95 with *Malat1* RNA revealed that *Malat1* was not colocalized with glutamatergic synapses (Supplemental Fig. 3E-H). Thus, the *Malat1* in neuronal processes is packaged with Staufen proteins into structures similar to mRNA granules.

Neuronal mRNAs traveling within dendrites can be mobilized for translation by synaptic stimulation, which induces their local unpackaging from the granule and increases their FISH signal (Holt et al. 2019; Kiebler and Bassell 2006; Schuman 1999; Krichevsky and Kosik 2001; Formicola et al. 2021; Buxbaum et al. 2014). Similarly, we found that depolarization of the cultured neurons with 60 mM potassium chloride for 1 hour led to a significant increase in the cytoplasmic FISH staining for *Malat1* (Fig. 3E-F). This increased signal was not due to an increase in *Malat1* RNA abundance as measured by RT/PCR (Supplemental Fig. 2G-I). Taken together these data indicate that cytoplasmic *Malat1* is localized to RNA granules in neuronal processes and is released in an activity-dependent manner.

### Small polypeptides are encoded within the 5’ region of *Malat1*

Many RNAs originally classified as noncoding have been found to encode small peptides serving a variety of cellular functions (Nelson et al. 2016; Matsumoto et al. 2017; Huang et al. 2017). The activity-dependent release of cytoplasmic *Malat1* from mRNA granules in neurons raised the possibility that *Malat1* might engage with ribosomes and be translated. To assess this, we examined ribosome profiling data of mRNAs in cultured cortical neurons. We identified several ribosome peaks within the 5’ region of *Malat1*, suggesting that *Malat1* associates with ribosomes (Fig. 4A). This was in contrast to two other lncRNAs, *Neat1* and *Norad*, that showed no ribosomal binding peaks (Supplemental Fig. 4B-C). Reexamining previously reported ribosome profiling data for dendritically localized mRNAs in rat neurons showed similar peaks to mouse *Malat1* (Saini et al. 2019). Similar peaks were also previously observed in human *Malat1* from HeLa cells (Wilusz et al. 2012). For typical neuronal mRNAs such as *Map2* (Supplemental Fig. 4A) ribosome occupancy was limited to the open reading frames. Searching for possible ORFs near the ribosomal peaks within *Malat1*, we identified 6 potential short ORFs (M1-M6), each with an ATG start codon and a minimal ORF length of 30 nt (Fig. 4A). These ORFs exhibited relatively modest but statistically significant conservation between mammalian species (Fig. 4B and Supplemental Fig. 5A-B). Of these ORFs only M1 showed overlapping ribosome binding and this did not extend equally through the ORF. Similar incomplete coverage has been observed in other short ORF’s (Powers et al. 2022; Brar et al. 2012). This ribosome association with *Malat1* could result in productive translation, serve some regulatory role, or simply be adventitious.

**Fig. 4:**
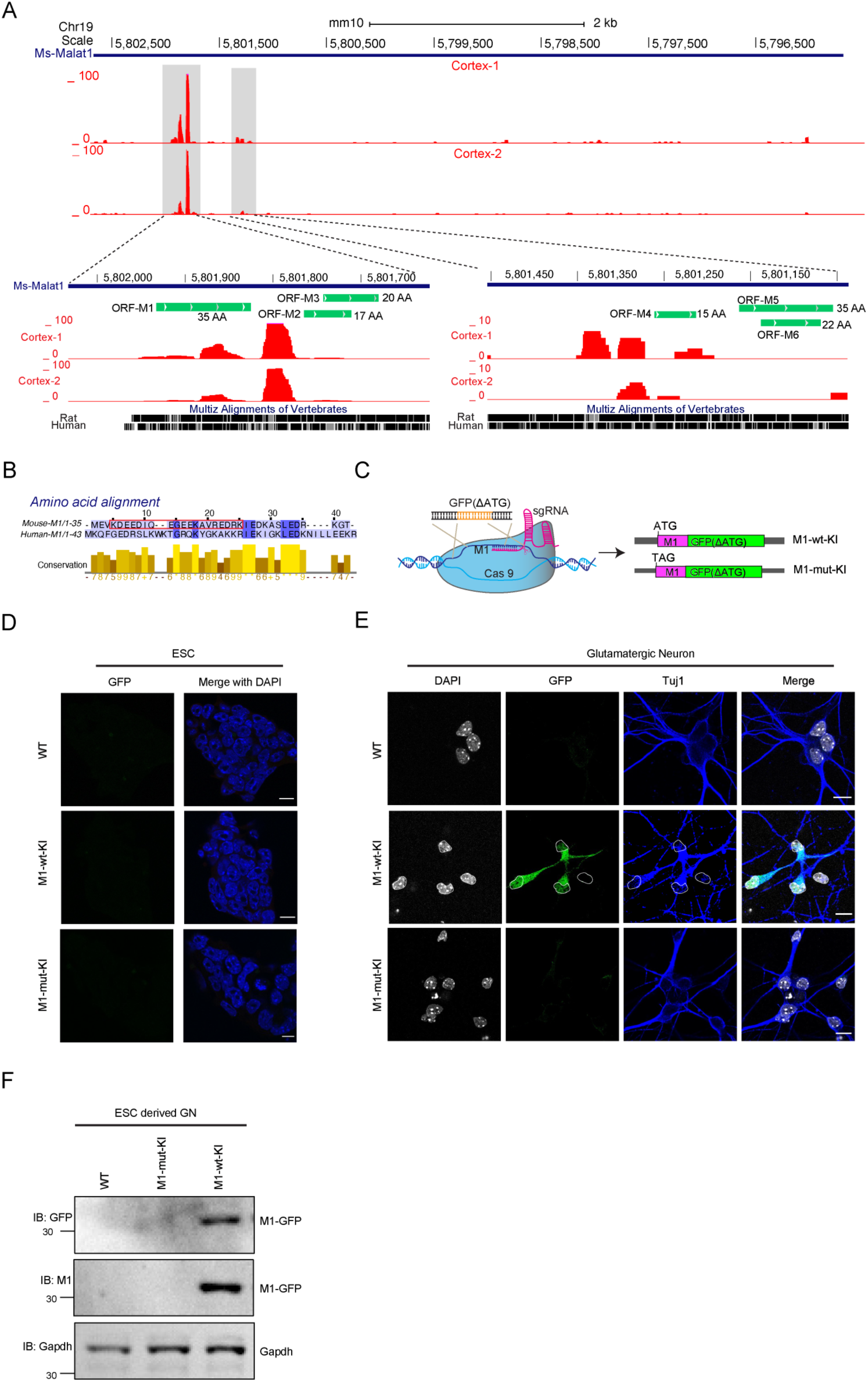
*Malat1* is bound by ribosomes and translated in neurons. (A) Genome browser view of the *Malat1* locus displaying ribosome profiling data in cultured cortical neurons. Lower panel, enlarged views of the grey highlighted ribosome binding peaks. Potential ORFs are shown as green bars with their peptide lengths. (B) Amino acid alignment of the mouse and human M1 peptides by clustal-W. The red box highlights the peptide sequence used as antigen to raise antibodies to the M1 protein. (C) Diagram of Crispr/Cas9 targeting to knock in the GFP coding sequence as a C-terminal fusion with M1 at the endogenous *Malat1* loci of embryonic stem cells. M1-WT-KI contains the M1 ATG initiation codon, and M1-Mut-KI has this codon mutated to TAG. (D) Fluorescent images of parental and genome edited ES cells showing a lack of GFP fluorescence in all three genotypes. (E) GFP fluorescence of WT and genome edited cells after differentiation into glutamatergic neurons. The knock-in cells containing the M1 initiation codon are now expressing GFP. (F) Immunoblot analysis of M1-GFP in glutamatergic neurons derived from engineered ESC lines probed with GFP and M1 antibodies.

To determine whether the *Malat1* ORFs are translated in neurons, we created fusion constructs containing the EGFP ORF, minus its own ATG, linked as an in-frame C-terminal extension of each *Malat1* ORF (Supplemental Fig. 6A). Transient expression of these constructs in N2a neuroblastoma cells and assay by fluorescence microscopy revealed that the M1 and M5 ORFs can initiate productive translation to produce the fused GFP (Supplemental Fig. 6B). Immunoblots confirmed that these proteins migrated at the expected masses of the M1 and M5 fusion proteins (Supplemental Fig. 6C). Notably, GFP expression was lost when the M1 start codon was mutated from ATG to TAG, indicating initiation is occurring at the M1 ATG (Supplemental Fig. 6A-C). To confirm that GFP was translated from the entire *Malat1* and not a fragment of the ectopically expressed RNA, we performed RNA FISH in N2a cells after transient expression of the M1-GFP and M1-mut-GFP constructs (Supplemental Fig. 6D). *Malat1* FISH signals were observed in both nucleus and cytoplasm (Supplemental Fig. 6D). The cytoplasmic signal for *Malat1* colocalized with the FISH signal for GFP, indicating that the GFP protein was translated from the full length *Malat1*. The M1-mutant-GFP-*Malat1* transcripts were localized to the cytoplasm, but did not produce GFP protein (Supplemental Fig. 6D). These results demonstrate that the M1 ORF is translated from the whole *Malat1* transcript when expressed from a plasmid.

### The M1 peptide is translated from RNA produced from the endogenous *Malat1* loci

To confirm that M1 peptide is produced from endogenous *Malat1*, we employed CRISPR-Cas9 editing of the *Malat1* locus in mouse ES cells (E14) to insert GFP as a C-terminal extension of the M1 ORF. This generated a knockin GFP-tagged M1 ORF (M1-wt-KI, Fig. 4C). As a negative control, a parallel construct replaced the M1 ATG start codon with TAG to create a mutant knockin allele of the M1 ORF (M1-mut-KI, Fig. 4C). Genotyping individual edited clones identified one homozygous and four heterozygous M1-wt-KI lines, as well as three heterozygous M1-mut-KI lines (Supplemental Fig. 7A-B). The correct in-frame insertion of GFP into the *Malat1* loci was confirmed by Sanger sequencing (Supplemental Fig. 7C). Neither the M1-wt-KI nor M1-mut-KI alleles exhibited GFP fluorescence in ES cells, as expected from the nuclear localization of *Malat1* (Fig. 4D). We then differentiated the wildtype and GFP-knock-in ES lines into glutamatergic neurons (Supplemental Fig. 7D). All three lines differentiated efficiently into cells with neuronal morphology that expressed neuronal markers Tuj1, PSD95, and vGlut1 as assayed by RT/PCR and immunofluorescence (Supplemental Fig. 7E-G). We found that GFP protein was expressed in the M1-wt-KI neurons but was absent in the M1-mut-KI neurons, indicating translation of the endogenous *Malat1* M1-ORF in differentiated neurons (Fig. 4E). The M1-GFP expression was selective to neurons and absent from non-neuronal cells in the culture. The expression of M1-GFP protein was also validated by immunoblot using GFP and M1 antibodies (described below) in the ESC-derived neurons (Fig. 4F). These data demonstrate that in neurons, but not in ESC, *Malat1* RNA undergoes translation initiation at the start codon of the M1 ORF.

### M1 peptide expression is enhanced by depolarization

To assay the presence of the M1 peptide without a GFP fusion, we raised an antibody to a 19 amino acid segment of the 35 residue peptide (Fig. 4B). We confirmed the reactivity and specificity of the antibody in immunoblot assays in N2a cells expressing an RFP-M1 fusion protein (Supplemental Fig. 8A-B). The M1 antibody also immunoprecipitated the GFP-M1 fusion protein (Supplemental Fig. 8B-C). In immunofluorescence assays of N2a cells expressing RFP fusion proteins, the antibody yielded abundant cytoplasmic staining in cells expressing RFP-M1 and no signal in cells expressing RFP fused to the M6 peptide or to the Rbfox1 protein (Supplemental Fig. 8D-E). These experiments confirmed that the M1 antibody could detect the protein with minimal background. The short length of the native M1 peptide precluded its detection by immunoblot.

Immunofluorescent staining of cultured neurons with the M1 antibody detected expression of the native protein in the cytoplasm and dendritic processes (Fig. 5A). To confirm that the fluorescent staining was derived from the M1 peptide and not another reactivity of the antibody, we treated the neurons with the Gapmer oligos to degrade *Malat1* (Supplemental Fig. 2D). Importantly, depletion of Malat1 eliminated the immunofluorescence staining by the M1 antibody (Fig. 5A). Thus, endogenous M1 peptide encoded by *Malat1* is produced in cultured neurons.

**Fig. 5:**
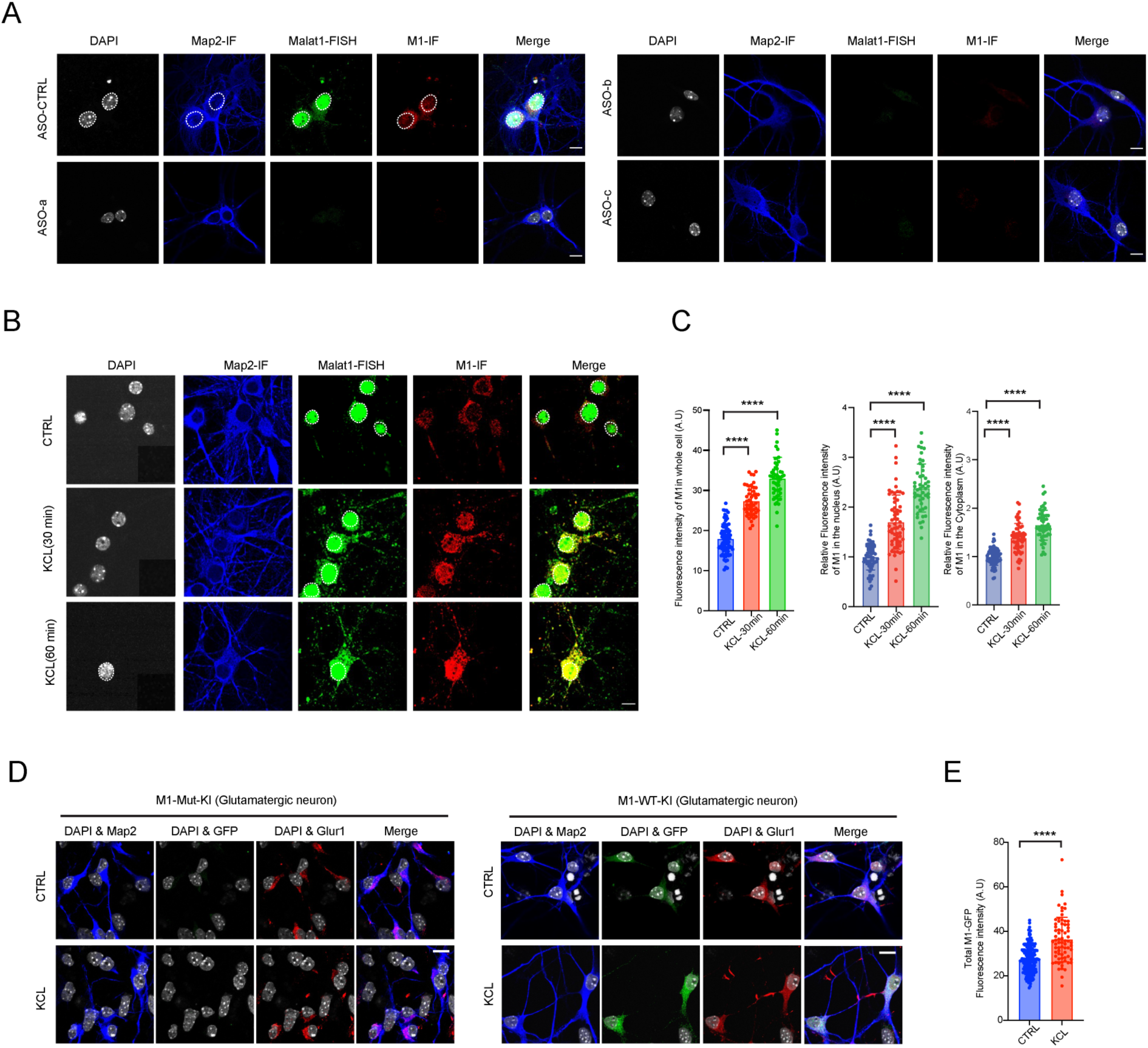
The M1 peptide is detected in primary neurons and is upregulated by excitatory stimuli. (A) M1 peptide immunofluorescence before and after *Malat1* knockdown by ASOs in primary neurons. (B) Antibody staining of the endogenous M1 peptide after KCL treatment for 30 min or 60 min in primary cortical neurons. (C) Left, quantification of mean IF intensities of 50 neuronal cells from b. Middle, relative mean fluorescence intensity of nuclear M1. Right, relative mean fluorescence intensity of cytoplasmic M1 (D) Immunofluorescence of Map2, M1-GFP and GluR1 in glutamatergic neurons derived from M1-wt-KI and M1-mut-KI ESC lines. GFP indicates signal for endogenous M1-GFP. Anti-Glur1 was used as a marker for glutamatergic neurons. (E) Quantification of mean fluorescent intensities for > 50 cells from d. “****” indicates P value < 0.0001. “NS” indicates P > 0.05. Scale bar, 10 um.

We showed above that KCl depolarization resulted in increased *Malat1* FISH signal in neurons without an increase in Malat1 abundance, presumably due to unpackaging of the RNA from neuronal granules (Krichevsky and Kosik 2001; Formicola et al. 2021; Buxbaum et al. 2014; Bi et al. 2006; Mofatteh et al. 2020). (Fig. 3E-F). To test if this activity-dependent release of local *Malat1* transcripts resulted in increased M1 micropeptide translation, we depolarized primary cortical neurons with 60 mM KCl (Ueda et al. 2022). We found that KCl depolarization for 30 and 60 min led to 1.5 and 2 fold increases respectively in M1 staining over the whole cell. The nuclear M1 protein was notably increased indicating that the small M1 peptide can enter the nucleus after synthesis (Fig. 5C). We also examined expression of M1-GFP in the ESC derived neurons. In these cells, GFP fluorescence increased 35% after KCL treatment (Fig. 5D-E). Overall, these data indicate that neurons translate *Malat1* RNA to produce the M1 peptide and that M1 peptide expression is increased with neuronal activity.

## Discussion

### *Malat1* is exported to the cytoplasm in neurons

We demonstrate that in postmitotic neurons a portion of the typically nuclear lncRNA *Malat1* is exported into the cytoplasm, where it is translated to produce a micropeptide (M1). The unusual processing pathway of *Malat1* and its lack of a poly-A tail do not preclude its translation (Wilusz et al. 2008, 2012; Brown et al. 2012). The 3’ portion of the *Malat1* RNA was previously shown to enhance translation of an upstream ORF when present in a reporter mRNA (Wilusz et al. 2012).

*Malat1* has also been found in the cytoplasm of several types of cancer cells, including bladder, hepatic, and breast cancer, and of platelet precursor cells (Zhu et al. 2023; Zhao et al. 2021; Shih et al. 2021; Sun et al. 2023). It is not clear if Malat1 RNA is translated in these cells, but it was found to encode an antigenic peptide in colorectal cancer cells (Barczak et al. 2023). The mechanisms that allow selective release of Malat1 from the nucleus are not clear. *Malat1* is normally sequestered on chromatin through a mechanism that requires binding by the U1 snRNP (Yin et al. 2020). In addition to splicing, U1 also functions to suppress premature polyadenylation during transcription (Venters et al. 2019). Interestingly, changes in U1 activity both after neuronal stimulation and in cancer cells are thought to cause the global activation of new cleavage and polyadenylation sites (Berg et al. 2012). Thus, the release of *Malat1* for nuclear export selectively in neurons may involve reduced activity or availability of U1 in these cells. However, U1 inhibition was found to release Malat1 from chromatin into the soluble nucleoplasm but not its export to the cytoplasm. Thus, changes in U1 function alone are unlikely to be sufficient for *Malat1* export. *Malat1* enrichment in nuclear speckles also requires multiple RNA binding proteins (Miyagawa et al. 2012; Wang et al. 2019). Inhibition of U1 or depletion of the nuclear speckle factors do not release Malat-1 into the cytoplasm. Additional factors mediating its nuclear localization could include an expression and nuclear retention element (ENE) similar to those found on viral noncoding RNAs (Brown et al. 2012; Conrad and Steitz 2005), and/or m6A modifications seen in synaptically localized *Malat1* (Madugalle et al. 2023).

### *Malat1* is a localized mRNA

We find that *Malat1* is packaged into neuronal granules that contain Staufen protein and are trafficked into neuronal processes of developing cortical neurons. Malat1 has been observed in neurites of hippocampal neurons by expansion microscopy (Alon et al. 2021), and was found to enrich in synaptic fractions after a fear extinction learning protocol (Madugalle et al. 2023). Dendritic RNA granules contain mRNAs that are translationally silent and masked to detection by FISH (Bauer et al. 2023; Buxbaum et al. 2014). They are transported along processes through association with microtubule based motors to allow their selective unpackaging and translational activation at specific stimulated synapses (Holt et al. 2019; Fritzsche et al. 2013). Similar to localized mRNA, both protease treatment and depolarization with KCl dramatically increase the detection of neuritic *Malat1* by FISH. Neuronal depolarization with KCl also increases synthesis of the *Malat1* encoded M1 peptide. These data together uncover a new function for *Malat1* as not only a nuclear lncRNA, but also a cytoplasmic coding RNA.

### Functions of *Malat1* translation products

We found that the peptide encoded by the M1 ORF is expressed from the endogenous *Malat1* locus in stimulated neurons. So far, the M1 peptide is the only *Malat1* translation product directly observed in neurons. We did observe modest translation of an M5 ORF-GFP fusion produced from a transgene in N2a cells. Peptides from additional *Malat1* ORFs may be synthesized in other cells or conditions. Further work interrogating the function of M1 and perhaps other peptides should shed light on the roles of micropeptides in neuronal maturation (Duffy et al. 2022).

The existence of cytoplasmic, translated *Malat1* must now be considered in interpreting the effects of *Malat1* depletion experiments. An earlier study found that loss of *Malat1* reduced expression of synaptic proteins in hippocampal neurons (Bernard et al. 2010). Others found that Malat-1 knockdown in N2a cells or hippocampal neurons inhibited neurite outgrowth (Chen et al. 2016; Jiang et al. 2020), whereas Malat-1 depletion from the brain was seen to impair fear-extinction memory (Madugalle et al. 2023). These observations in diverse settings could all involve loss of the M1 peptide along with the RNA. We found that depletion of *Malat1* from neurons, using either ASO’s or shRNAs (data not shown), stimulated the expression of synaptic and other neuronal proteins. These divergent observations from those earlier (Bernard et al. 2010) could arise from differences in cell types, culture systems, or methods of modulating *Malat1* levels and will need further investigation. In our system, the stimulated expression of synaptic proteins observed upon *Malat1* depletion could result from the loss of nuclear *Malat1* RNA, as proposed in earlier studies, or from the loss of the M1 peptide. It is also possible that loss of *Malat1* from the pool of ribosome bound RNAs might have an indirect effect on translational capacity.

The presence of *Malat1* as a translating mRNA in neurons also suggests an new possible source for physiological phenotypes observed in the *Malat1* knockout mice. These mice develop normally and phenotypes from Malat1 loss have primarily been observed in either the nervous system or in cancer. It will be interesting to assess during the late neuroendocrine state of many cancers whether Malat1 becomes cytoplasmic and produces M1 peptide. The role of micropeptides in these cellular processes will be an interesting area to explore.

## SUPPLEMENTAL INFORMATION

Supplemental Information including nine figures and two tables and can be found with this article.

## MATERIALS AND METHODS

### Tissue culture

We maintained mouse embryonic stem cells (E14) in ESC Media containing: DMEM (Fisher Scientific) supplemented with 15 % ESC-qualified fetal bovine serum (Thermo Fisher Scientific), 1x non-essential amino acids (Thermo Fisher Scientific), 1x GlutaMAX (Thermo Fisher Scientific), 1x ESC-qualified nucleosides (EMD Millipore), 0.1 mM β-Mercaptoethanol (Sigma-Aldrich), and 10^3^ units/ml ESGRO leukemia inhibitor factor (LIF) (EMD Millipore). N2a cells were maintained in Dulbecco’s Modified Eagle’s Medium (DMEM) (Gibco, Invitrogen) supplemented with 10% fetal bovine serum (FBS) and penicillin-streptomycin. Cells were grown in incubator with 5% CO_2_ at 37C.

### Primary cortical neuron culture

Embryonic day 16 C57BL/6J pregnant dams (Charles River Laboratories) were sacrificed by CO_2_ overdose followed by cervical dislocation. Embryos were decapitated with sharp scissors, and cortices from males and females were dissected into ice-cold Hank’s Balanced Salt Solution (HBSS, Ca2+- and Mg2+-free) and randomly pooled. Cortices were treated with DNase1 and trypsin in a 37C water bath for 12 min. Cortices were then washed once with HBSS and triturated in HBSS containing 10% DNase1 by pipetting up and down for 12 times. Dissociated cells were spun down and washed once with Plating Media (Neurobasal supplemented with 20% horse serum, 10% 250 mM sucrose in neurobasal, 0.25x Glutamax and 1x Pen/Strep). Cortical neurons were plated at a density of ∼500 cells/mm^2^ (for RNA or protein isolation) or ∼250 cells/mm^2^ (for immunocytochemistry) on tissue culture plates or coverslips (Fisher Scientific, NC0672873) coated with 0.1mg/mL poly-L-lysine (Sigma-Aldrich, P1274-100mg) in borate buffer (0.1 M borate acid in H_2_O, pH8.5). Cells were initially plated in Plating Media and then refreshed with Feeding Media (Neurobasal supplemented with B27 and Glutamax) the second day after seeding. AraC was added at DIV4 to a final concentration of 2.5 uM. Half the culture media was replaced with fresh Feeding Media every 3 days beginning at 4 days in vitro (DIV4). Primary cultures were maintained in a 37C incubator supplemented with 5% CO_2_.

### GapmeR ASO Knockdown

Cortical primary neurons were isolated from E16 embryos and plated at a density of ∼250 cells/mm2 on poly-L-lysine coated plates or coverslips. GapmeR ASOs were gymnotically introduced into primary neurons at DIV 8. GapmeRs were synthesized by IDT and transfected into cells as previously described (Williams et al. 2022). Briefly, For gymnotic delivery, the ASO was added to the medium at the desired concentration (typically 2.5 to 5 uM) with a single treatment at DIV 8. ASO’s were not replenished with fresh medium additions. After 3 days transfection, the cells were harvested for RNA extraction or immunofluorescence. The Control and Malat1 knockdown ASOs are list below.

Control-ASO sequences:

<u>5′-/52MOErC/*/i2MOErC/*/i2MOErT/* /i2MOErT/*C*C* C*T*G* A*A*G* G*T*T* C*/i2MOErC/*/i2MOErT/* /i2MOErC/*/32MOErC/-3’</u>

Malat1-ASO-a sequences:

<u>5′-/52MOErG/*/i2MOErG/*/i2MOErG/*/i2MOErT/*/i2MOErC/*A*G*C*T*G*C*C*A*A*T*/ i2MOErG/*/i2MOErC/*/i2MOErT/*/i2MOErA/*/32MOErG/ −3′</u>

Malat1-ASO-b sequences:

<u>5′-/52MOErC/*/i2MOErC/*/i2MOErA/* /i2MOErG/*G*C* T*G*G* T*T*A* T*G*A* C*/i2MOErT/*/i2MOErC/* /i2MOErA/*/32MOErG/ −3′</u>

Malat1-ASO-c sequences:

<u>5’-/52MOErA/*/i2MOErA/*/i2MOErC/* /i2MOErT/*A*C* C*A*G* C*A*A* T*T*C*/i2MOErC/*/i2MOErG/*/i2MOErC/* /32MOErC/ - 3’.</u>

### Ribosome profiling

Primary cortical neurons were dissected from E16 embryos in C57BL/6 mice. The dissociated neurons were then plated on 10 cm dishes at a density of 2.25 million cells. The primary neurons were cultured for 18 days. Cells were flash frozen in liquid nitrogen at DIV 18, moved to dry ice, and lysed in Turbo DNase I lysis buffer containing 100 ug/ml cycloheximide. Cell lysate was digested with RNase I at a ratio of 1U RNase:2 ug RNA for 45 minutes on a nutator. The reaction was inhibited with Superase-In RNase Inhibitor (Thermo Fisher Scientific catalog number AM 2696). Ribosome protected fragments (RPFs) were pelleted through a sucrose cushion centrifuged for 2 h at 100,0000xg in a TLA centrifuge at 4 degrees Celsius. Ribosome protected RNA fragments were recovered using the Zymo Direct RNA Miniprep kit. RNA was precipitated with isopropanol and resuspended in 10 mM Tris PH 8. Footprint fragments were purified by gel electrophoresis on a 15% polyacrylamide TBE-Urea gel stained with SYBR Gold. A 10 bp ladder, NI-800, and NEB miRNA were used as markers to select and isolate 17-34 nt fragments ∼28nt footprints. RNA was extracted from gel slices overnight, precipitated with isopropanol, and resuspended in 10 mM Tris pH8. Footprints were then dephosphorylated and ligated to pre-adenylated 3’ linkers with unique barcodes. Ligation reactions were purified using the Zymo Oligo Clean and Concentrator Kits. Next, ribosomal RNA (rRNA) was depleted using RiboZero Gold Illumina kit according to manufacturer’s protocol. Samples were again purified with the Clean and Concentrator Kit. Reverse transcription was performed, and cDNA was circularized using circligase II and library was constructed using PCR. Distribution analysis was conducted using HSD1000 Screen Tape and verified to be on average between 175-190 bp in length. Libraries were sequenced and aligned to whole mouse genome. All steps were conducted using two biological replicates.

### Immunofluorescence (IF)

ESC, N2A and cultured primary neuron cells were washed once with ice-cold PBSM (1 x PBS, 5 mM MgCl_2_), followed by fixation with 4% paraformaldehyde in PBSM for 10 min at room temperature (RT). After a 5-minute wash with ice-cold PBSM, the cells were permeabilized with 0.3% Triton X-100 in PBSM for 7 min on ice. The cells were washed once with PBSM and blocked with 3% BSA (Fraction V) in PBSM for 0.5 h at RT. The coverslips were then incubated with primary antibody in 3% BSA in PBSM for 1 h at RT. After 3 washes in PBST (1xPBS 0.1% tween 20), secondary antibody (goat-anti-mouse-Cy3: VWR-95040-042 or goat-anti-rabbit-cy5 or Donkey anti-chicken-488) diluted in in 1 x PBS was added for 45 min at RT.

For IF only: Cells were washed three times with PBST and then stained with DAPI in PBST for 15 min. Cells were mounted with prolong mounting media overnight at room temperature overnight. Antibodies used in this study. MAP2 (Abcam, ab5392), STAU1(Abcam, ab73478), STAU2 (Thermo, PA5-78473) Synaptophysin (sysy-101004), PSD95 (Antibodies Incorporated, 75-028), GFP (Abcam, ab290), Tuj1 (Abcam, ab18207), M1 antibody generated from Thermo Scientific (Project 1XJ0541), Vglut1 (Synaptic Systems, 135 303), Glur1 (Thermo, MA5-27694).

For IF combined with RNA FISH : After 3 washes in 1x PBST, cells were refixed in 4% paraformaldehyde in PBSM for 10 min at RT. After a brief wash with PBSM, cells were equilibrated in 10% formamide in 2 x SSC for 30 min. FISH probes were hybridized to cells at a concentration of 0.5 ng/ul in Hybridization buffer (Biosearch: SMF-HB1-10, 10% formamide added freshly) on parafilm, and placed in a humidified box overnight. Cells were washed once with Wash-buffer A (Biosearch: SMF-WA1-60) at 37C for 30 min followed by washing with Wash-buffer A containing 0.5 ug/ml DAPI at 37C for 30 min. Cells were washed once with Wash-buffer B (Biosearch: SMF-WB1-20) at RT for 5 min and mounted with prolong anti-fade mounting media until completely dry. Slides were subject for confocal microscopy.

## ACKNOWLEDGMENTS

We thank Dr. Prasanth Kannanganattu for *Malat1* plasmids, Dr. David Spector for advice and materials, members of the D.L.B. lab for helpful discussions, the UCLA Neuroscience Genomics Core, and the imaging core of the California Nanosystems Institute at UCLA. This work was supported by NIH grant R35GM136426, a research grant from the Broad Stem Cell Research Center at UCLA, and a research grant from the WM Keck Foundation to DLB.

## AUTHOR CONTRIBUTIONS

Concept and experimental design, W.X. and D.L.B.; Experiment execution, W.X., R.H. and K-H.Y.; Experimental materials M.N.; Sequence data processing, C-H.L.; Analysis of experimental data, W.X. and D.L.B.; Writing manuscript, W.X. and D.L.B.; Review & Editing, all authors.

## REFERENCES

Adam SA. 2016. Nuclear Protein Transport in Digitonin Permeabilized Cells. In The Nuclear Envelope (eds. S. Shackleton, P. Collas, and E.C. Schirmer), Vol. 1411 of *Methods in Molecular Biology*, pp. 479–487, Springer New York, New York, NY http://link.springer.com/10.1007/978-1-4939-3530-7_29 (Accessed October 10, 2023).

Alon S, Goodwin DR, Sinha A, Wassie AT, Chen F, Daugharthy ER, Bando Y, Kajita A, Xue AG, Marrett K, et al. 2021. Expansion sequencing: Spatially precise in situ transcriptomics in intact biological systems. Science 371: eaax2656.

Anderson DM, Anderson KM, Chang C-L, Makarewich CA, Nelson BR, McAnally JR, Kasaragod P, Shelton JM, Liou J, Bassel-Duby R, et al. 2015. A Micropeptide Encoded by a Putative Long Noncoding RNA Regulates Muscle Performance. Cell 160: 595–606.

Barczak W, Carr SM, Liu G, Munro S, Nicastri A, Lee LN, Hutchings C, Ternette N, Klenerman P, Kanapin A, et al. 2023. Long non-coding RNA-derived peptides are immunogenic and drive a potent anti-tumour response. Nat Commun 14: 1078.

Batish M, Van Den Bogaard P, Kramer FR, Tyagi S. 2012. Neuronal mRNAs travel singly into dendrites. Proc Natl Acad Sci USA 109: 4645–4650.

Bauer KE, De Queiroz BR, Kiebler MA, Besse F. 2023. RNA granules in neuronal plasticity and disease. Trends in Neurosciences 46: 525–538.

Berg MG, Singh LN, Younis I, Liu Q, Pinto AM, Kaida D, Zhang Z, Cho S, Sherrill-Mix S, Wan L, et al. 2012. U1 snRNP Determines mRNA Length and Regulates Isoform Expression. Cell 150: 53–64.

Bergmann JH, Spector DL. 2014. Long non-coding RNAs: modulators of nuclear structure and function. Current Opinion in Cell Biology 26: 10–18.

Bernard D, Prasanth KV, Tripathi V, Colasse S, Nakamura T, Xuan Z, Zhang MQ, Sedel F, Jourdren L, Coulpier F, et al. 2010. A long nuclear-retained non-coding RNA regulates synaptogenesis by modulating gene expression. EMBO J 29: 3082–3093.

Bi J, Tsai N-P, Lin Y-P, Loh HH, Wei L-N. 2006. Axonal mRNA transport and localized translational regulation of κ-opioid receptor in primary neurons of dorsal root ganglia. Proc Natl Acad Sci USA 103: 19919–19924.

Bi P, Ramirez-Martinez A, Li H, Cannavino J, McAnally JR, Shelton JM, Sánchez-Ortiz E, Bassel-Duby R, Olson EN. 2017. Control of muscle formation by the fusogenic micropeptide myomixer. Science 356: 323–327.

Böhmdorfer G, Wierzbicki AT. 2015. Control of Chromatin Structure by Long Noncoding RNA. Trends in Cell Biology 25: 623–632.

Brar GA, Yassour M, Friedman N, Regev A, Ingolia NT, Weissman JS. 2012. High-Resolution View of the Yeast Meiotic Program Revealed by Ribosome Profiling. Science 335: 552–557.

Brown JA, Valenstein ML, Yario TA, Tycowski KT, Steitz JA. 2012. Formation of triple-helical structures by the 3ʹ-end sequences of MALAT1 and MENβ noncoding RNAs. Proc Natl Acad Sci USA 109: 19202–19207.

Buxbaum AR, Wu B, Singer RH. 2014. Single β-Actin mRNA Detection in Neurons Reveals a Mechanism for Regulating Its Translatability. Science 343: 419–422.

Chen L, Feng P, Zhu X, He S, Duan J, Zhou D. 2016. Long non-coding RNA Malat1 promotes neurite outgrowth through activation of ERK / MAPK signalling pathway in N2a cells. J Cell Mol Med 20: 2102–2110.

Chen X, He L, Zhao Y, Li Y, Zhang S, Sun K, So K, Chen F, Zhou L, Lu L, et al. 2017. Malat1 regulates myogenic differentiation and muscle regeneration through modulating MyoD transcriptional activity. Cell Discov 3: 17002.

Conrad NK, Steitz JA. 2005. A Kaposi’s sarcoma virus RNA element that increases the nuclear abundance of intronless transcripts. EMBO J 24: 1831–1841.

Duffy EE, Finander B, Choi G, Carter AC, Pritisanac I, Alam A, Luria V, Karger A, Phu W, Sherman MA, et al. 2022. Developmental dynamics of RNA translation in the human brain. Nat Neurosci 25: 1353–1365.

Engreitz JM, Sirokman K, McDonel P, Shishkin AA, Surka C, Russell P, Grossman SR, Chow AY, Guttman M, Lander ES. 2014. RNA-RNA Interactions Enable Specific Targeting of Noncoding RNAs to Nascent Pre-mRNAs and Chromatin Sites. Cell 159: 188–199.

Formicola N, Heim M, Dufourt J, Lancelot A-S, Nakamura A, Lagha M, Besse F. 2021. Tyramine induces dynamic RNP granule remodeling and translation activation in the Drosophila brain. eLife 10: e65742.

Fritzsche R, Karra D, Bennett KL, Ang F yee, Heraud-Farlow JE, Tolino M, Doyle M, Bauer KE, Thomas S, Planyavsky M, et al. 2013. Interactome of Two Diverse RNA Granules Links mRNA Localization to Translational Repression in Neurons. Cell Reports 5: 1749–1762.

Grzejda D, Mach J, Schweizer JA, Hummel B, Rezansoff AM, Eggenhofer F, Panhale A, Lalioti M- E, Cabezas Wallscheid N, Backofen R, et al. 2022. The long noncoding RNA *mimi* scaffolds neuronal granules to maintain nervous system maturity. Sci Adv 8: eabo5578.

Gueroussov S, Gonatopoulos-Pournatzis T, Irimia M, Raj B, Lin Z-Y, Gingras A-C, Blencowe BJ. 2015. An alternative splicing event amplifies evolutionary differences between vertebrates. Science 349: 868–873.

Holt CE, Martin KC, Schuman EM. 2019. Local translation in neurons: visualization and function. Nat Struct Mol Biol 26: 557–566.

Huang J-Z, Chen M, Chen D, Gao X-C, Zhu S, Huang H, Hu M, Zhu H, Yan G-R. 2017. A Peptide Encoded by a Putative lncRNA HOXB-AS3 Suppresses Colon Cancer Growth. Molecular Cell 68: 171–184.e6.

Ingolia NT, Brar GA, Stern-Ginossar N, Harris MS, Talhouarne GJS, Jackson SE, Wills MR, Weissman JS. 2014. Ribosome Profiling Reveals Pervasive Translation Outside of Annotated Protein-Coding Genes. Cell Reports 8: 1365–1379.

Jiang T, Cai Z, Ji Z, Zou J, Liang Z, Zhang G, Liang Y, Lin H, Tan M. 2020. The lncRNA MALAT1/miR-30/Spastin Axis Regulates Hippocampal Neurite Outgrowth. Front Cell Neurosci 14: 555747.

Karakas D, Ozpolat B. 2021. The Role of LncRNAs in Translation. ncRNA 7: 16.

Khanduja JS, Calvo IA, Joh RI, Hill IT, Motamedi M. 2016. Nuclear Noncoding RNAs and Genome Stability. Molecular Cell 63: 7–20.

Kiebler MA, Bassell GJ. 2006. Neuronal RNA Granules: Movers and Makers. Neuron 51: 685– 690.

Kim J, Piao H-L, Kim B-J, Yao F, Han Z, Wang Y, Xiao Z, Siverly AN, Lawhon SE, Ton BN, et al. 2018. Long noncoding RNA MALAT1 suppresses breast cancer metastasis. Nat Genet 50: 1705–1715.

Knowles RB, Sabry JH, Martone ME, Deerinck TJ, Ellisman MH, Bassell GJ, Kosik KS. 1996. Translocation of RNA Granules in Living Neurons. J Neurosci 16: 7812–7820.

Krichevsky AM, Kosik KS. 2001. Neuronal RNA Granules. Neuron 32: 683–696.

Lee S, Kopp F, Chang T-C, Sataluri A, Chen B, Sivakumar S, Yu H, Xie Y, Mendell JT. 2016. Noncoding RNA NORAD Regulates Genomic Stability by Sequestering PUMILIO Proteins. Cell 164: 69–80.

Madugalle SU, Liau W-S, Zhao Q, Li X, Gong H, Marshall PR, Periyakaruppiah A, Zajaczkowski EL, Leighton LJ, Ren H, et al. 2023. Synapse-Enriched m ^6^ A-Modified Malat1 Interacts with the Novel m ^6^ A Reader, DPYSL2, and Is Required for Fear-Extinction Memory. J Neurosci 43: 7084–7100.

Mallardo M, Deitinghoff A, Müller J, Goetze B, Macchi P, Peters C, Kiebler MA. 2003. Isolation and characterization of Staufen-containing ribonucleoprotein particles from rat brain. Proc Natl Acad Sci USA 100: 2100–2105.

Matsumoto A, Pasut A, Matsumoto M, Yamashita R, Fung J, Monteleone E, Saghatelian A, Nakayama KI, Clohessy JG, Pandolfi PP. 2017. mTORC1 and muscle regeneration are regulated by the LINC00961-encoded SPAR polypeptide. Nature 541: 228–232.

Mattick JS, Amaral PP, Carninci P, Carpenter S, Chang HY, Chen L-L, Chen R, Dean C, Dinger ME, Fitzgerald KA, et al. 2023. Long non-coding RNAs: definitions, functions, challenges and recommendations. Nat Rev Mol Cell Biol 24: 430–447.

Miao H, Wu F, Li Y, Qin C, Zhao Y, Xie M, Dai H, Yao H, Cai H, Wang Q, et al. 2022. MALAT1 modulates alternative splicing by cooperating with the splicing factors PTBP1 and PSF. Sci Adv 8: eabq7289.

Miyagawa R, Tano K, Mizuno R, Nakamura Y, Ijiri K, Rakwal R, Shibato J, Masuo Y, Mayeda A, Hirose T, et al. 2012. Identification of *cis* - and *trans*-acting factors involved in the localization of MALAT-1 noncoding RNA to nuclear speckles. RNA 18: 738–751.

Mofatteh M, 2020. mRNA localization and local translation in neurons. AIMS Neuroscience 7: 299–310.

Munschauer M, Nguyen CT, Sirokman K, Hartigan CR, Hogstrom L, Engreitz JM, Ulirsch JC, Fulco CP, Subramanian V, Chen J, et al. 2018. The NORAD lncRNA assembles a topoisomerase complex critical for genome stability. Nature 561: 132–136.

Nakagawa S, Ip JY, Shioi G, Tripathi V, Zong X, Hirose T, Prasanth KV. 2012. Malat1 is not an essential component of nuclear speckles in mice. RNA 18: 1487–1499.

Nelson BR, Makarewich CA, Anderson DM, Winders BR, Troupes CD, Wu F, Reese AL, McAnally JR, Chen X, Kavalali ET, et al. 2016. A peptide encoded by a transcript annotated as long noncoding RNA enhances SERCA activity in muscle. Science 351: 271–275.

Niklas J, Melnyk A, Yuan Y, Heinzle E. 2011. Selective permeabilization for the high-throughput measurement of compartmented enzyme activities in mammalian cells. Analytical Biochemistry 416: 218–227.

Noh JH, Kim KM, McClusky WG, Abdelmohsen K, Gorospe M. 2018. Cytoplasmic functions of long noncoding RNAs. WIREs RNA 9. https://onlinelibrary.wiley.com/doi/10.1002/wrna.1471 (Accessed May 31, 2023).

Ouyang J, Zhong Y, Zhang Y, Yang L, Wu P, Hou X, Xiong F, Li X, Zhang S, Gong Z, et al. 2022. Long non-coding RNAs are involved in alternative splicing and promote cancer progression. Br J Cancer 126: 1113–1124.

Powers EN, Chan C, Doron-Mandel E, Llacsahuanga Allcca L, Kim Kim J, Jovanovic M, Brar GA. 2022. Bidirectional promoter activity from expression cassettes can drive off-target repression of neighboring gene translation. eLife 11: e81086.

Ransohoff JD, Wei Y, Khavari PA. 2018. The functions and unique features of long intergenic non-coding RNA. Nat Rev Mol Cell Biol 19: 143–157.

Ruiz-Orera J, Messeguer X, Subirana JA, Alba MM. 2014. Long non-coding RNAs as a source of new peptides. eLife 3: e03523.

Saini H, Bicknell AA, Eddy SR, Moore MJ. 2019. Free circular introns with an unusual branchpoint in neuronal projections. eLife 8: e47809.

Sato K, Sakai M, Ishii A, Maehata K, Takada Y, Yasuda K, Kotani T. 2022. Identification of embryonic RNA granules that act as sites of mRNA translation after changing their physical properties. iScience 25: 104344.

Schuman EM. 1999. mRNA Trafficking and Local Protein Synthesis at the Synapse. Neuron 23: 645–648.

Shih C-H, Chuang L-L, Tsai M-H, Chen L-H, Chuang EY, Lu T-P, Lai L-C. 2021. Hypoxia-Induced MALAT1 Promotes the Proliferation and Migration of Breast Cancer Cells by Sponging MiR-3064-5p. Front Oncol 11: 658151.

Sun Y, Wang T, Lv Y, Li J, Jiang X, Jiang J, Zhang D, Bian W, Zhang C. 2023. MALAT1 promotes platelet activity and thrombus formation through PI3k/Akt/GSK-3β signalling pathway. Stroke Vasc Neurol 8: 181–192.

Tang Y, Wang J, Lian Y, Fan C, Zhang P, Wu Y, Li X, Xiong F, Li X, Li G, et al. 2017. Linking long non-coding RNAs and SWI/SNF complexes to chromatin remodeling in cancer. Mol Cancer 16: 42.

Tripathi V, Ellis JD, Shen Z, Song DY, Pan Q, Watt AT, Freier SM, Bennett CF, Sharma A, Bubulya PA, et al. 2010. The Nuclear-Retained Noncoding RNA MALAT1 Regulates Alternative Splicing by Modulating SR Splicing Factor Phosphorylation. Molecular Cell 39: 925–938.

Ueda HH, Nagasawa Y, Sato A, Onda M, Murakoshi H. 2022. Chronic neuronal excitation leads to dual metaplasticity in the signaling for structural long-term potentiation. Cell Reports 38: 110153.

Venters CC, Oh J-M, Di C, So BR, Dreyfuss G. 2019. U1 snRNP Telescripting: Suppression of Premature Transcription Termination in Introns as a New Layer of Gene Regulation. Cold Spring Harb Perspect Biol 11: a032235.

Wang C, Lu T, Emanuel G, Babcock HP, Zhuang X. 2019. Imaging-based pooled CRISPR screening reveals regulators of lncRNA localization. Proc Natl Acad Sci USA 116: 10842–10851.

Wang H, Wang Y, Xie S, Liu Y, Xie Z. 2016. Global and cell-type specific properties of lincRNAs with ribosome occupancy. Nucleic Acids Res gkw909.

Williams LA, Gerber DJ, Elder A, Tseng WC, Baru V, Delaney-Busch N, Ambrosi C, Mahimkar G, Joshi V, Shah H, et al. 2022. Developing antisense oligonucleotides for a TECPR2 mutation-induced, ultra-rare neurological disorder using patient-derived cellular models. Molecular Therapy - Nucleic Acids 29: 189–203.

Wilusz JE, Freier SM, Spector DL. 2008. 3ʹ End Processing of a Long Nuclear-Retained Noncoding RNA Yields a tRNA-like Cytoplasmic RNA. Cell 135: 919–932.

Wilusz JE, JnBaptiste CK, Lu LY, Kuhn C-D, Joshua-Tor L, Sharp PA. 2012. A triple helix stabilizes the 3ʹ ends of long noncoding RNAs that lack poly(A) tails. Genes Dev 26: 2392–2407.

Xiao W, Yeom K-H, Lin C-H, Black DL. 2023. Improved enzymatic labeling of fluorescent in situ hybridization probes applied to the visualization of retained introns in cells. RNA 29: 1274–1287.

Xie S-J, Diao L-T, Cai N, Zhang L-T, Xiang S, Jia C-C, Qiu D-B, Liu C, Sun Y-J, Lei H, et al. 2021. mascRNA and its parent lncRNA MALAT1 promote proliferation and metastasis of hepatocellular carcinoma cells by activating ERK/MAPK signaling pathway. Cell Death Discov 7: 110.

Xing J, Liu H, Jiang W, Wang L. 2021. LncRNA-Encoded Peptide: Functions and Predicting Methods. Front Oncol 10: 622294.

Yeom K-H, Pan Z, Lin C-H, Lim HY, Xiao W, Xing Y, Black DL. 2021. Tracking pre-mRNA maturation across subcellular compartments identifies developmental gene regulation through intron retention and nuclear anchoring. Genome Res 31: 1106–1119.

Yin Y, Lu JY, Zhang X, Shao W, Xu Y, Li P, Hong Y, Cui L, Shan G, Tian B, et al. 2020. U1 snRNP regulates chromatin retention of noncoding RNAs. Nature 580: 147–150.

Young AP, Jackson DJ, Wyeth RC. 2020. A technical review and guide to RNA fluorescence in situ hybridization. PeerJ 8: e8806.

Zhang C, Zhou B, Gu F, Liu H, Wu H, Yao F, Zheng H, Fu H, Chong W, Cai S, et al. 2022. Micropeptide PACMP inhibition elicits synthetic lethal effects by decreasing CtIP and poly(ADP-ribosyl)ation. Molecular Cell 82: 1297–1312.e8.

Zhang Q, Vashisht AA, O’Rourke J, Corbel SY, Moran R, Romero A, Miraglia L, Zhang J, Durrant E, Schmedt C, et al. 2017. The microprotein Minion controls cell fusion and muscle formation. Nat Commun 8: 15664.

Zhao Y, Zhou L, Li H, Sun T, Wen X, Li X, Meng Y, Li Y, Liu M, Liu S, et al. 2021. Nuclear-Encoded lncRNA MALAT1 Epigenetically Controls Metabolic Reprogramming in HCC Cells through the Mitophagy Pathway. Molecular Therapy - Nucleic Acids 23: 264–276.

Zhu N, Ahmed M, Li Y, Liao JC, Wong PK. 2023. Long noncoding RNA MALAT1 is dynamically regulated in leader cells during collective cancer invasion. Proc Natl Acad Sci USA 120: e2305410120.

