## Supplemental figures, materials and methods for "The lncRNA *Malat1* is trafficked to the cytoplasm as a localized mRNA encoding a small peptide in neurons"

**This PDF file includes:**

SI Materials and Methods

Figures S1 to S8

SI References

### Supplemental Experimental Procedures

#### Plasmids.

pSV2-mMalat1-full-length plasmid was a kind gift from the Kannanganattu V. Prasanth lab. To insert GFP as a C-terminal in frame extension to the ORF's (M1, M3, M4, M5 and M6), we first added an Age1 restriction site (accggt) downstream of each ORF by primer extension PCR using the primers listed below.

| | |
|---|---|
| Age1-M1-for | accggtTAAAGGCACAGGTAGTAGGAAGCC |
| Age1-M1-rev | AGTACCTTTTCTGTCTTCTAAAGATGCTTTG |
| Age1-M3-for | accggtTAAAGTCAGAAGGAAATTATCTTTAAAGCCA |
| Age1-M3-rev | TATCTTATCCAACCTTTTGGCCTCAATCTTAT |
| Age1-M4-for | accggtTAGGAAGAAGGAGCCATACGGATG |
| Age1-M4-rev | GCTTCCTCCACCGAACCGCA |
| Age1-M5-for | accggtTAGATAGCGGCTCCTAGACCAG |
| Age1-M5-rev | TCCTCTCCACAATGCCTGCTC |
| Age1-M6-for | accggtTAGATAGCGGCTCCTAGACCAG |

The GFP fragment (without its own ATG) with the flanking portion of Malat1 sequences were amplified by primer extension PCR using the following primers.

| | |
|---|---|
| GFP-Age1-M1-for | CTTTAGAAGACAGAAAAGGTACTGTGAGCAAGGGCGAGGAGC |
| GFP-Age1-M1-rev | TTCCTACTACCTGTGCCTTTACTTGTACAGCTCGTCCATGCC |
| GFP-Age1-M3-for | GCCAAAAGGTTGGATAAGATAGTGAGCAAGGGCGAGGAGC |
| GFP-Age1-M3-rev | GATAATTTCCTTCTGACTTTACTTGTACAGCTCGTCCATGCC |
| GFP-Age1-M4-for | GTGCGGTTCGGTGGAGGAAGCGTGAGCAAGGGCGAGGAGC |
| GFP-Age1-M4-rev | CCGTATGGCTCCTTCTTCCTACTTGTACAGCTCGTCCATGCC |
| GFP-Age1-M5-for | GAGAGGATAGATAGCGGCTCCGTGAGCAAGGGCGAGGAGC |
| GFP-Age1-M5-rev | CACACTGGCATGCTGGTCTACTTGTACAGCTCGTCCATGCC |
| GFP-Age1-M6-for | GAGCAGGCATTGTGGAGAGGAGTGAGCAAGGGCGAGGAGC |
| GFP-Age1-M6-rev | TGGTCTAGGAGCCGCTATCTACTTGTACAGCTCGTCCATGCC |

The amplified GFP fragments and linearized vectors (by Age I) were ligated using the Gibson assembly kit (Thermo, A46627). The M1 mutant ORF (with the ATG mutated to TAG) plasmid was constructed using the primers below:

Primers:

| | |
|---|---|
| M1-ATG-mut-for | taGGAAGTGAAAGACGAAGAAGACA |

| | |
|---|---|
| M1-ATG-mut-rev | TTTCTAATCTTCTTTCTCCAAATACTAGCCTAAC |

For the RFP-M1 or RFP-M6 expression plasmids, M1 or M6 insert sequences were synthesized from IDT with Nhe1 and EcoR1 overhangs and then ligated in frame with downstream of RFP ORF in the pAAV-nEF vector. The RFP-Rbfox1 plasmid was gift from Celine Vuong.

**Genotyping analysis.**

The genomic DNAs of GFP knock-in ES lines were extracted using Quick-DNA Micro-prep Kit (Zymo Research, D3021). The inserted GFP in the Malat1 loci was validated by PCR using the primers below (See Supplemental Fig. 7a).

GFP internal primer:

GFP-FOR (F1): GTGAGCAAGGGCGAGGAG

GFP-REV (R1): TGTAGTTGTACTCCAGCTTGTGC

M1-flanking-for (F2): CCAGCCTGGTCTACAGAGTGA

M1-flanking-rev (R2): TGTGGCTGTCTAAAGACTCTTC.

The GFP insertion bands were excised and subjected to Sanger sequencing (see See Supplemental Fig. 7b).

**Crispr/Cas9 knock-in in ES cells.**

Both the M1-locus sgRNAs (sequence: TACTTAAAGGCACAGGTAGT) and spCas9 proteins were purchased from Synthego (CRISPR evolution sgRNA EZ Kit). The sgRNA was complementary to the sequence downstream of M1 ORF. M1 sgRNAs were dissolved in nuclease-free water to a concentration of 100 uM. The KI DNA templates (M1-wt-KI and M1-mut-KI) for homolog recombination were obtained by PCR and then column purified. The concentration of template was measured by Nanodrop before mixing with Cas9 RNP complex. The primers used for PCR amplification of the KI templates (wt and mut) are as listed below:

Malat1-M1-KI-For: 5'- CCAGCCTGGTCTACAGAGTGA-3'

Malat1-M1-KI-Rev: 5'- TGTGGCTGTCTAAAGACTCTTC-3'.

The RNP complex (spCas9 protein, sgRNAs) and the M1-KI-template were mixed in resuspension R buffer as follows:

| Component | Volume |
|---|---|

| | |
|---|---|
| KI-template(ssDNA) | 1.5 ul (20 pmol) |
| Resuspension buffer | 3.5 µl |
| sgRNA | 1 µl (100 pmol) |
| Cas9 | 1 µl (20 pmol) |
| Total volume | 7 µl |

The RNPs were incubated for 10 min at RT. ES cells were then trypsinized and dissociated. 1 million ES cells were suspended in 5 ul resuspension R buffer and mixed with 7 ul RNP mixture. A Neon electroporator was used to transfect RNP and DNA templates into cells. The electroporation conditions were used as follows:

pulse voltage: 1250 v,

pulse width: 20 ms,

pulse number: 2.

After electroporation, the mixture was added into growth media without serum. Cells were cultured for a week and genomic DNAs were extracted for genotyping analysis.

For the positive pools, the cells were serially diluted in 96- well format to approximately one cell per well. Monoclones were then dissociated and seeded into 24 well plates. After 4-5 days growth, cells were passaged to 6-well plates. Half of cells were subjected to genomic extraction. The other half were cultured for maintenance.

**Digitonin fractionation.**

For neuronal fraction at different DIVs, in vitro cultured neurons were washed once with 1 x PBS on a 10 cm dish. After the PBS was removed, 1 ml digitonin (20 uM) was gently added to the dish and incubated at RT for 4 min with agitation. After digitonin permeabilization, the digitonin buffer was transferred to a new tube as cytoplasmic fraction. The remaining cells on the plate were washed once with 5 ml 1 x PBS and then lysed with 1ml RIPA buffer. The lysate from RIPA buffer was defined as cytoplasmic-nuclear fraction. RNAs were extracted from the cytoplasmic and cytoplasmic-nuclear fractions by TRIzol and subjected for RT-qPCR.

For neuronal fractionation with/without KCL treatment, primary neurons were cultured in one well of 6-well plates. After washing cells once with 1xPBS, 150 ul of 20 uM digitonin was added to cells with 4 min agitation. After permeabilization, the digitonin buffer was transferred to a new tube as cytoplasmic fraction. 500 ul tryzol was added and mixed well with the cytoplasmic fraction. The left over cells on the well were washed once with 1xPBS. PBS was removed and another 500 ul tryzol was added into cells as cytoplasmic-nuclear fraction. RNA was extracted by RNeasy Mini Kit (74106) from Qiagen.

The qPCR primer sequences are listed in Table S1.

**Quantitative real-time PCR.**

RNA was isolated using TRIzol and reversed transcribed into cDNA with random hexamers using Superscript III Reverse Transcriptase (Thermo, 18-080-085). qRT-PCR was performed in 384-well plates using Sensi-FAST SYBR Lo-ROX Kit (Bioline, BIO-94020) for detection on a Quant studio 6 flex thermal cycler (Applied Biosystems). Relative gene expression was obtained using CT values of Gapdh or RPL32 for normalization. The qPCR primer sequences are listed in Table S1.

**Immunoblotting.**

In vitro cultured neuronal cells was lysed in RIPA buffer (150 mM sodium chloride, 50 mM Tris-HCl, 1 mM EDTA, 1% nonidet P40, 0.1% SDS, 0.5% Sodium deoxycholate ) with protease and phosphatase inhibitors (Roche, 5056489001) and benzonase (Sigma Aldrich, E1014-25KU). Lysates were centrifuged, cleared, and boiled in NuPAGE 1x LDS sample buffer (Thermo, NP0007), separated on 4- 2% Bis-Tris Protein Gels (NP0335BOX) and transferred to 0.45 mm PVDF membranes (GE Amersham) or NC membrane (Fisher Scientific, PI88018). Membranes were imaged on an iBright FL1500 Imaging System. Primary antibodies used in this study were: GFP (Abcam, ab290), Gapdh (Proteintech, 60004-1-IG), RFP (thermos, MA5-15257), M1 antibody generated from Thermo Scientific (Project 1XJ0541), Synaptophysin (sysy-101004), PSD95 (Antibodies Incorporated, 75-028), STAU2 (Proteintech, 15998-1-AP), Tuj1 (Abcam, ab18207).

**RNA FISH.**

RNA FISH probes were house made and labeled with distinct dyes (Xiao et al. 2023). Malat1 FISH oligos were ordered from IDT in a wet 96-well format. The FISH protocol was previously described (Xiao et al. 2023). Briefly, ESC, N2a and cultured primary neuron cells were washed once with ice-cold PBSM (1xPBS, 5mM $MgCl_2$), followed by fixation with 4% paraformaldehyde in PBSM for 10 min at room temperature (RT). After a 5-minute wash with ice-cold PBSM, the cells were permeabilized with 70% ethanol overnight. The cells were hydrated with 2 x SSC buffer containing 10% formamide for 1 h. RNA FISH probes were hybridized to cells at a concentration of 0.5 ng/ul in Hybridization buffer (Biosearch: SMF-HB1-10, 10% formamide added freshly). Coverslips were placed in a humidified box overnight. Cells were washed once with Wash-buffer A (Biosearch: SMF-WA1-60) at 37C for 30 min followed by washing with Wash-buffer A containing 0.5 ug/ml DAPI at 37C for another 30 min. Cells were washed once with Wash-buffer B (Biosearch: SMF-WB1-20) or 2 x SSC buffer at RT for 5 min and mounted with 8 ul prolong anti-fade mounting media overnight at RT. The RNA FISH probe sequences for Malat1, Gapdh and Camk2a are list in Table S2.

**RNA FISH assay after Proteinase K treatment.**

Cultured neurons were washed with 1 x PBS and then fixed with 4% PFA in PBS for 15 minutes at RT. Cells were soaked in 70% ethanol overnight at 4C. After ethanol permeabilization, cells were washed 2 times with 2 x SSC buffer. 20 mg/ml proteinase K was diluted 1000 times in 2 x SSC buffer to final concentraton of 20 ug/ml and then added to the fixed cells for 15 min at 37C. Cells were rinsed with 2 x SSC and fixed with 4% PFA in PBS again for 5 min. Cells were then washed once with 2 x SSC with 10% formamide. FISH probes in hybridization buffer were then added and the rest of steps were the same as FISH protocols described above.

**Differentiation of mouse ESCs to GNs.**

E14 ESCs were differentiated to glutamatergic neurons (GNs) as previously described (Gueroussov et al. 2015) with some modifications. Briefly, feeder-free mouse ESCs were grown as aggregate culture for eight days (DIV-8) in differentiation media consisting of Glasgow's MEM (GMEM) (Thermo Fisher Scientific) supplemented with 5% Knockout Serum Replacement (KSR) (Invitrogen), non-essential amino acids (Life Technologies), 1mM Sodium Pyruvate (Invitrogen), GlutaMAX (Life Technologies), and 0.1 mM 2-Mercaptoethanol (Sigma-Aldrich). Differentiating aggregates/embryoid bodies (EBs) were maintained at 37°C, 5% $CO_2$ and 90% relative humidity, with 50% of the media replaced by fresh media in every two days. After four days, the differentiation media was supplemented with 5 µM all-trans retinoic acid (RA, Sigma-Aldrich) until day 8. On day 8 (DIV 0), EBs/aggregates were dissociated with TrypLE Express Enzyme (Thermo Fisher Scientific) for 10 minutes at room temperature and dissociation was stopped by Trypsin inhibitor from Glycine max (Sigma-Aldrich). Single cells were then passed through a 40 µm nylon strainer (Thermo Fisher Scientific) and plated in N-2 media consisting of Neurobasal-A medium (Invitrogen) supplemented with N-2 Supplement (Life Technologies), GlutaMAX (Life Technologies), and 100 U/mL Penicillin-Streptomycin (Thermo Fisher Scientific) on Poly-D-lysine hydrobromide (Sigma-Aldrich) and Laminin (Sigma-Aldrich) coated dishes. Complete media changes were done 4 hours and 24 hours after plating. Two days later (DIV 2), N-2 media was replaced by B-27 media consisting of B-27 Serum-Free Supplement (Thermo Fisher Scientific), GlutaMAX (Life Technologies), and 100 U/mL Penicillin-Streptomycin (Thermo Fisher Scientific). Differentiating neurons were refreshed with new media every 3 days. On DIV-5, the media was supplemented with 10 µM 5'-fluoro-2'-deoxyuridine (Sigma-Aldrich) and 30 µM Uridine (Sigma-Aldrich) to inhibit remaining glia until DIV-10.

**Neuronal depolarization.**

Primary cortical neurons were cultured in vitro for 9 days, KCl was added in the media to a final concentration of 60 mM for 30 min or 60 min. Equal amout of $H_2O$ was added as negative control. For ESC derived Glutamatergic neuron treated with KCl, 60 mM KCl was added for 1 h, cells were then fixed with 4% PFA for immunofluorescence.

**Quantification of Pre- and Post- synaptic Puncta**

All image quantification was done manually in FIJI (Version: 2.3.0/1.53f). Cultured neurons used for quantification were imaged on a Leica sp8 light sheet confocal microscope in a Z-stack at 63x magnification. For cytoplasmic Malat1 puncta quantification, punctal numbers per 10 micrometers along the processes were counted manually from 2 cultures (biological replicates). For colocalization with pre- and post-synaptic markers, Synaptophysin or PSD95 intensities were measured along neurites and axons starting from the soma. The length of neurites was set and measured according to MAP2 staining. We set the width of segmented line to 40 pixels. The length of area of interest was set to 30 microns away from the soma. The mean intensity within this area of interest was measured according to those settings for all the neurons.

For the mean intensity of M1 expression, a segmented line with the width of 40 pixels was drawn along the neurites. The segmented line started from the soma and extended across neurites for 30 um (constant area for every neuron). The mean gray value was measured and plotted accordingly.

**Ribosome profiling analysis.**

Sequenced reads were demultiplexed using HtSeqTools. Input sequence reads linkers and adaptors were removed with cutadapt. Trimmed reads shorter than 24nt were discarded. Reads were then aligned to mm10 rRNA sequences with bowtie in default parameters and unaligned reads were retained. Retained reads were then aligned to the mm10 genome using STAR 2.5.4. Ribosome profiling data for cultured cortical neurons (Fig. 4a, Supplemental Fig. 4) is available in the GEO under accession number GSE249095.

**Statistical analysis**

Statistical analyses were performed in Prism 9. Data are represented by the mean, and error bars represent SD. Statistical tests performed are unpaired two-tailed t test. In the figures "*" indicates P value ≤ 0.05, "**" indicates P value ≤ 0.01, "***" indicates P value ≤ 0.001, "****" indicates P value < 0.0001, "NS" indicates P > 0.05.

**Supplemental Fig. 1**: Malat1 is transported to the cytoplasm in mouse and rat primary neurons during neuron development.

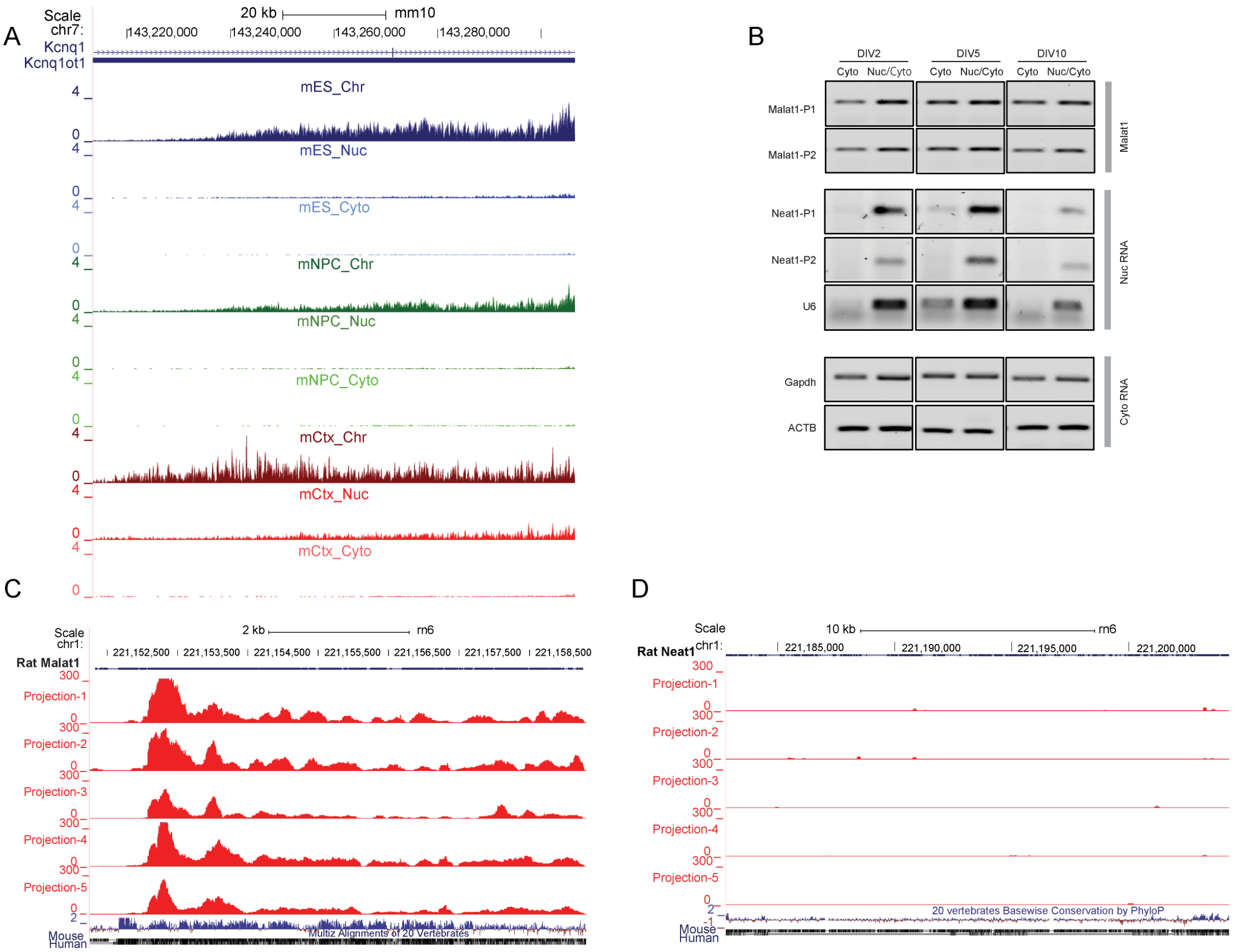


(A) Genome browser view of lncRNA Kcnq1ot1 expression in the chromatin, nucleoplasm, and cytoplasm fractions of three cell types. Blue, mouse embryonic cells; Green, mouse neuronal progenitor cells; Red, mouse primary cortical neurons (DIV 5). (B) RT-PCR analysis of *Malat1*, the nuclear noncoding RNAs *Neat1* and *U6*, and the cytoplasmic mRNAs *Gapdh* and *ACTB* after digitonin fractionation of cultured primary cortical neurons at different DIVs. P1 and P2 stand for primer pairs 1 and 2 targeting different regions of the gene of interest. PCR was performed with 25 cycles. (C) and (D) Genome browser tracks of *Malat1* and *Neat1* expression in neuronal projections from rat cultured hippocampal neurons. Data was downloaded at GSE129924 as generated by reference 4 (Saini et al. 2019).

**Supplemental Fig.2**: Validation of the specificity of Malat1 RNA FISH and knockdown, and quantification of Malat1 after KCl treatment.

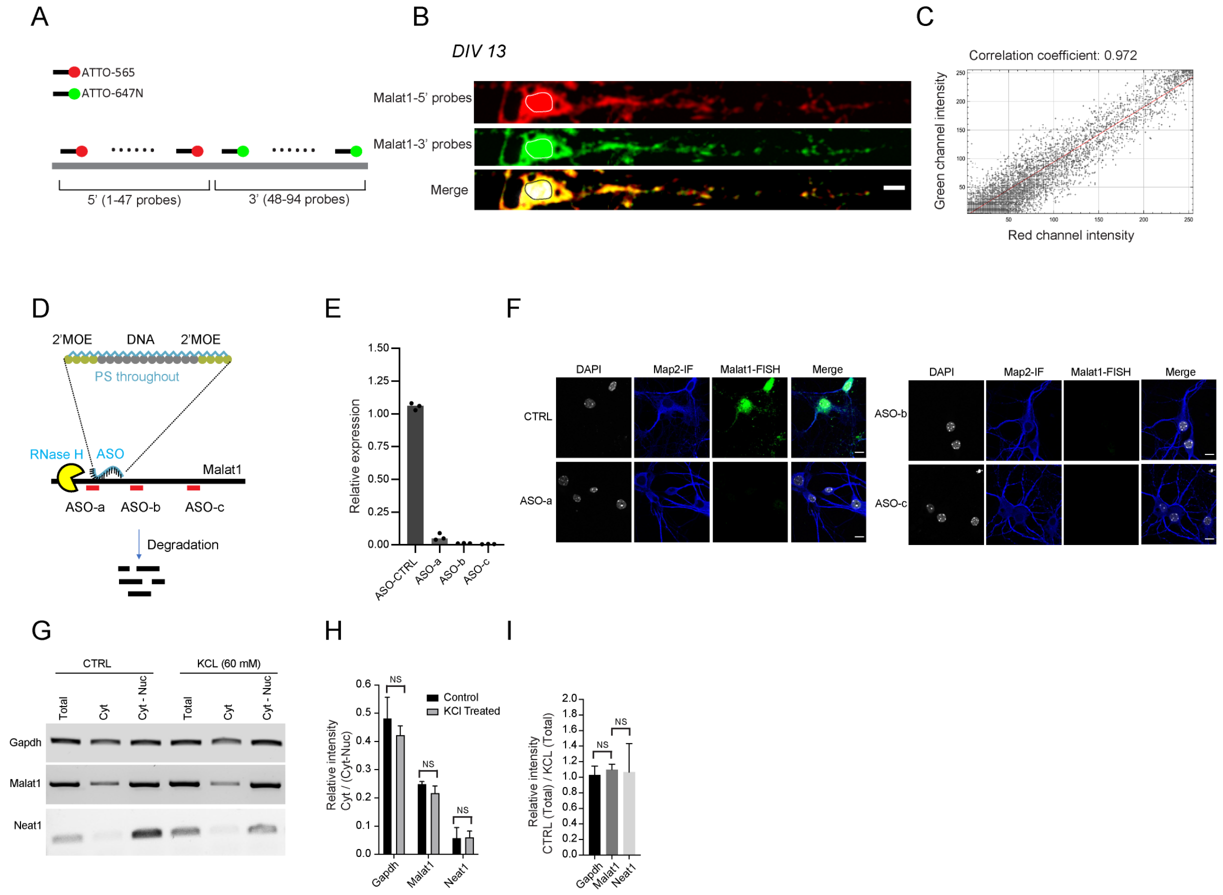


(A) Diagram of two sets of FISH probes targeting the 5’ and 3’ regions of mouse Malat1. Red probes (1 - 47) were labeled with ATTO-565 and green probes (48 - 94) with ATTO647N dye. (B) RNA FISH for Malat1 using distinct fluorescent probes (shown in a) in primary neurons at DIV13. Scale bar, 5 um. (C) Pixel intensity correlation of the 5’ and 3’ *Malat1* signals. (D) Diagram of Gapmer ASOs designed to induce RNase H of *Malat1*. ASO targeting sites within *Malat1* are indicated by three red bars. (E) qPCR analysis of Malat1 knockdown efficiency by Gapmer ASOs. (F) FISH analysis for validation of Malat1 Gapmer ASOs. (G) RT-PCR analysis of Malat1, Neat1 and Gapdh expression in total, cytosol, cytosol-nuclear fractions in control and KCL treated primary neurons at DIV9. The cells were fractionated by digitonin (see method). (H) quantification of the relative band intensities as the ratio of cyto / cyto-nuclear RNA for three genes from (G). (I)

quantification of the total band intensities of three genes in control and KCL treated cells shown in (G).

**Supplemental Fig.3**: co-localization of *Malat1* with proteins and RNAs.

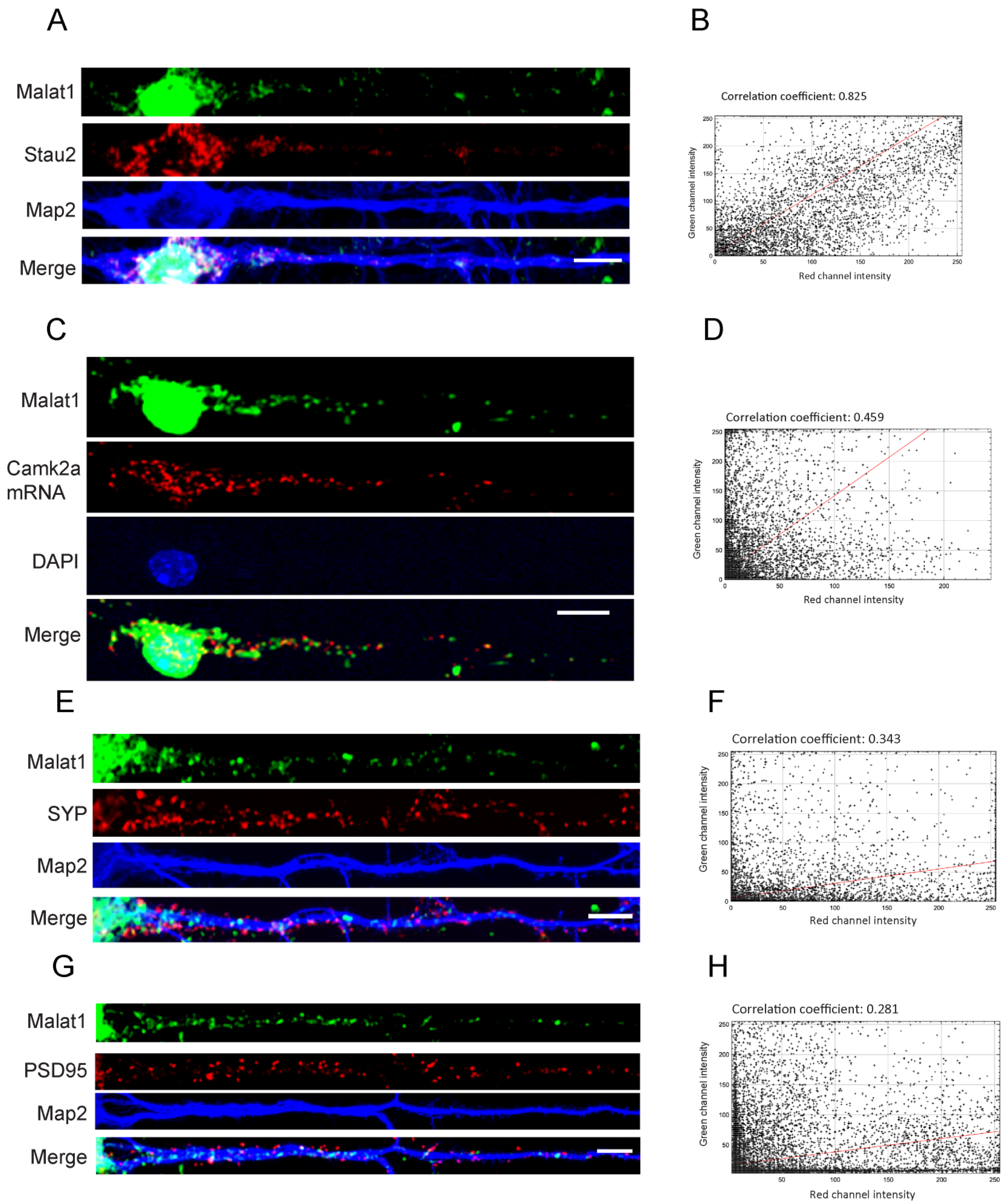

(A) *Malat1* RNA FISH combined with Map2 and Stau2 protein staining in cultured cortical neurons at DIV 13. (B) Pixel intensity correlation of the Stau2 and *Malat1* signals. (C) FISH of *Malat1* and *Camk2a* mRNA after proteinase K treatment in primary neurons at DIV13. (D) Pixel intensity correlation of the *Camk2a* and *Malat1* signals. (E) *Malat1* RNA FISH combined with immunofluorescence of Synaptophysin (SYP) and Map2 protein in cultured cortical neurons at DIV13. (F) Pixel intensity correlation of the *SYP* and *Malat1* signals. (G) *Malat1* RNA FISH combined with immunofluorescence of PSD95 and Map2 protein in cultured cortical neurons at DIV13. (H) Pixel intensity correlation of the PSD95 and *Malat1* signals. Scale bar, 5 um.

**Supplemental Fig.4**: Ribosomal footprints of coding and noncoding genes.

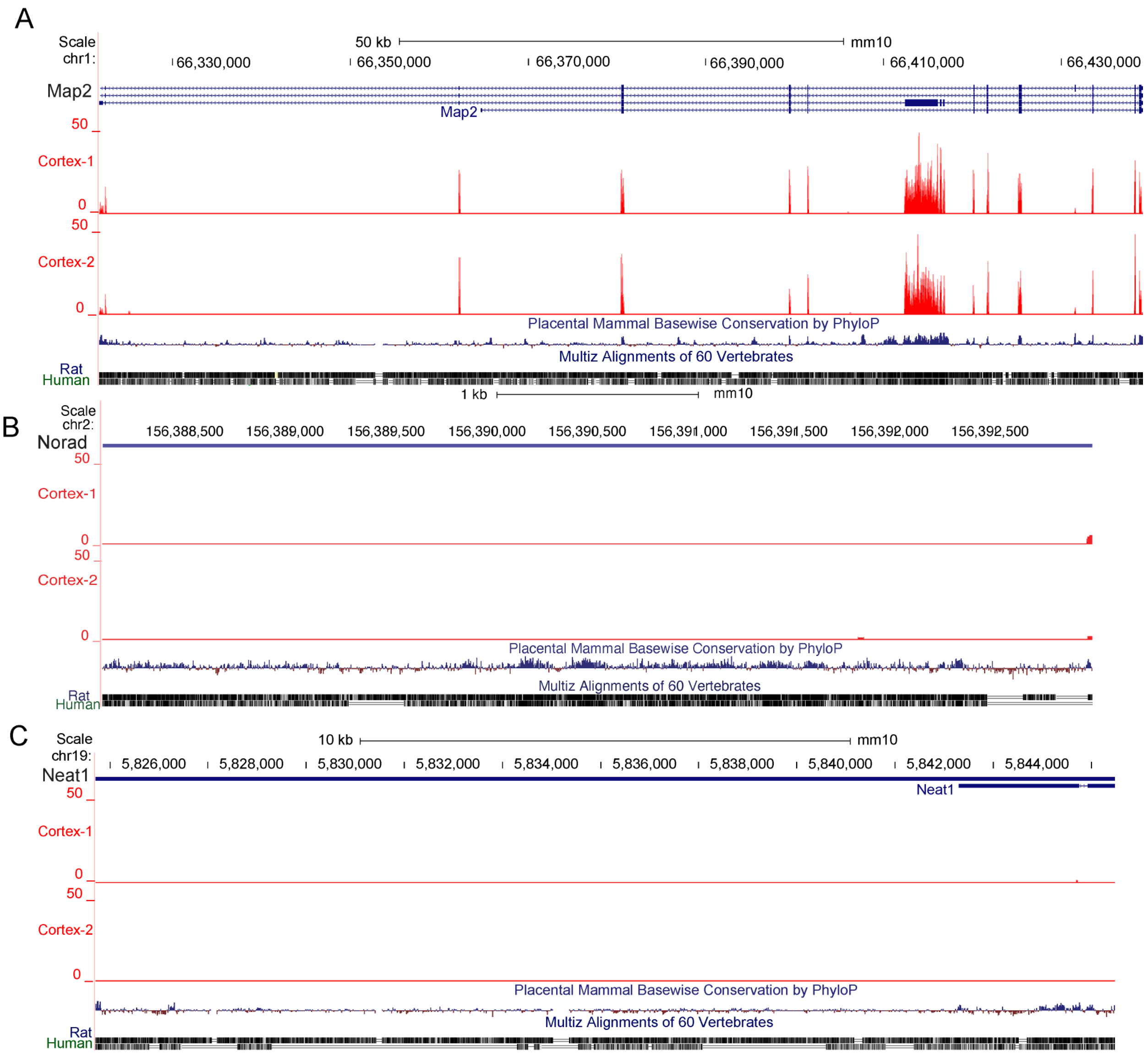


(A-C) Genome browser tracks for the ribosome binding peaks of an mRNA (*Map2*) and two lncRNAs (*Norad* and *Neat1*) in cultured mouse cortical neurons.

**Supplemental Fig.5**: Sequences of Malat1 ORFs across species.

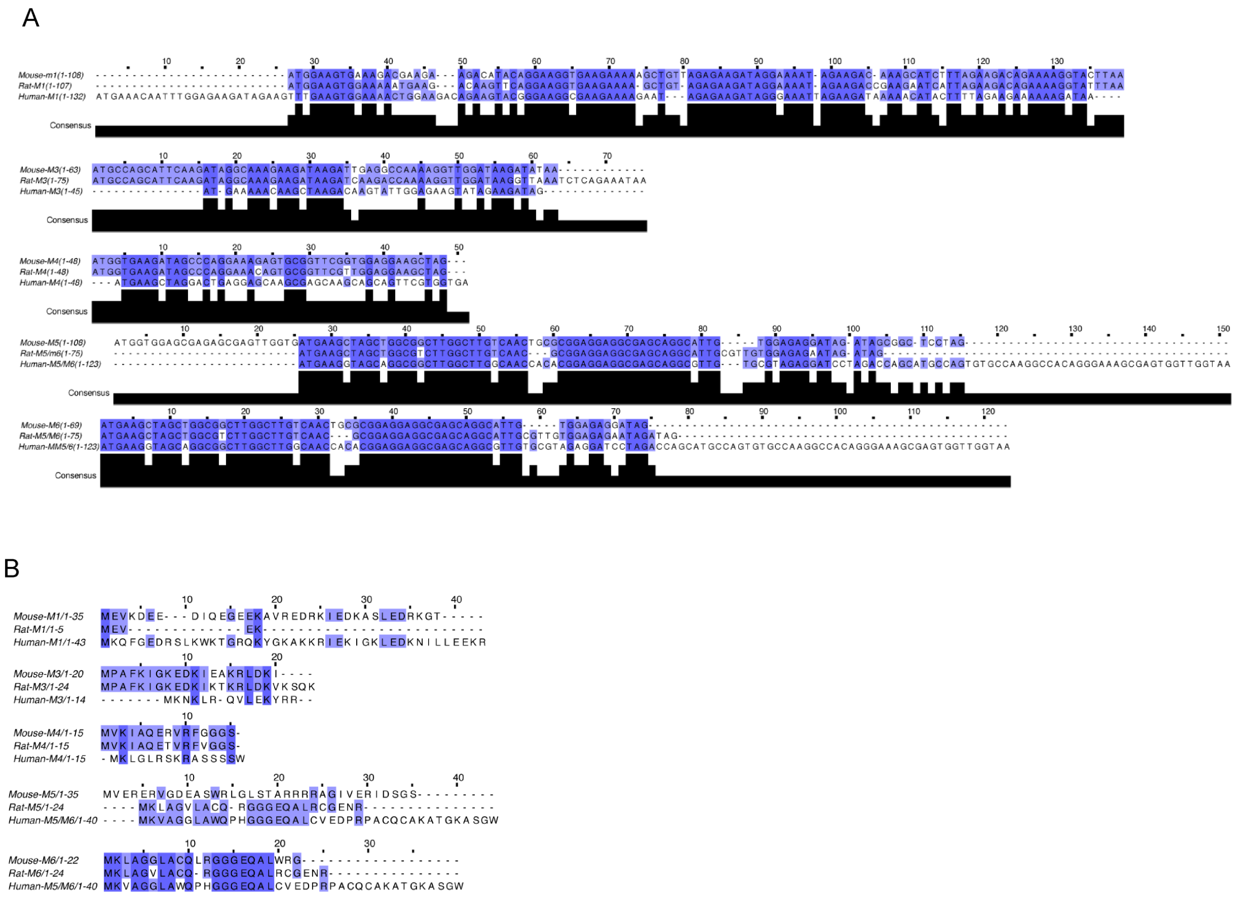


(A) Nucleotide sequences of the *Malat1* ORFs from mouse, rat and human aligned by Clustal-W2. (B) Amino acid sequences of the *Malat1* ORFs (M1, M3, M4, M5 and M6) from mouse, rat and human aligned by Clustal-W2. The M2 ORF did not exhibit conservation of either nucleotide or amino acid sequence.

**Supplemental Fig.6**: Malat1 is exported to the cytoplasm and encodes small peptides in N2a cells.

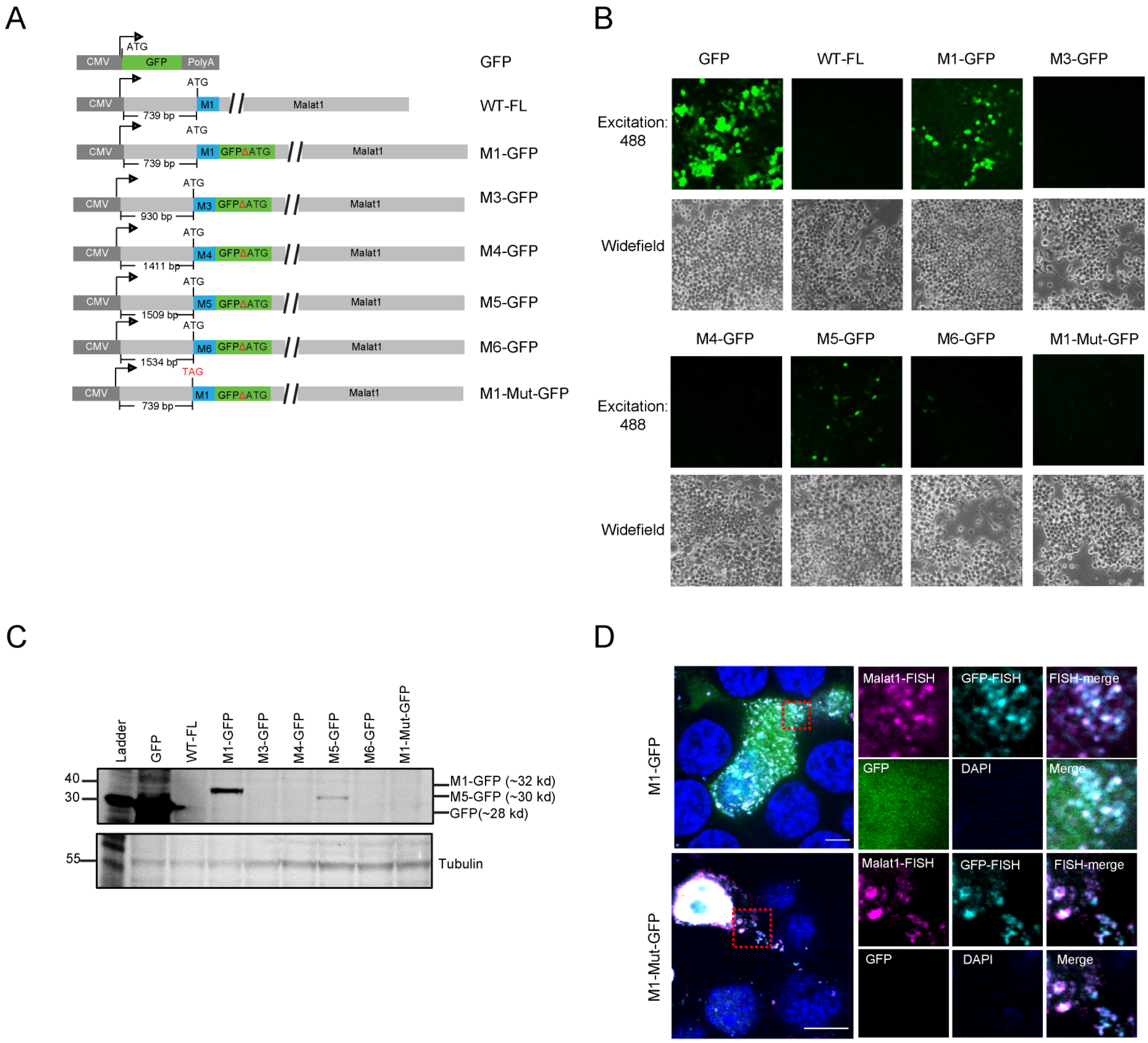


(A) Schematic of GFP and *Malat1* constructs with GFP inserted in-frame to different ORFs. The GFP plasmid served as a positive control. Full length *Malat1* plasmid WT-FL serves as a negative control. GFP sequence with its start codon deleted was appended in-frame to each ORF of the *Malat1* transcript. For the M1-mut-GFP plasmid, the M1 start codon ATG was mutated to TAG. (B) GFP fluorescence visualized after transient expression of the constructs in N2a cells (See A). (C) Western blot analysis of transiently expressed ORF-GFP constructs in N2a cells. Expected sizes of GFP-fusion proteins were labeled on the right. (D) *Malat1* and GFP RNA FISH, and the GFP protein fluorescence signal are shown in N2a cells transiently expressing the M1-WT-GFP and M1-Mut-GFP constructs.

**Supplemental Fig.7**: M1-GFP fusion protein is expressed from the endogenous Malat1 locus in glutamatergic neurons.

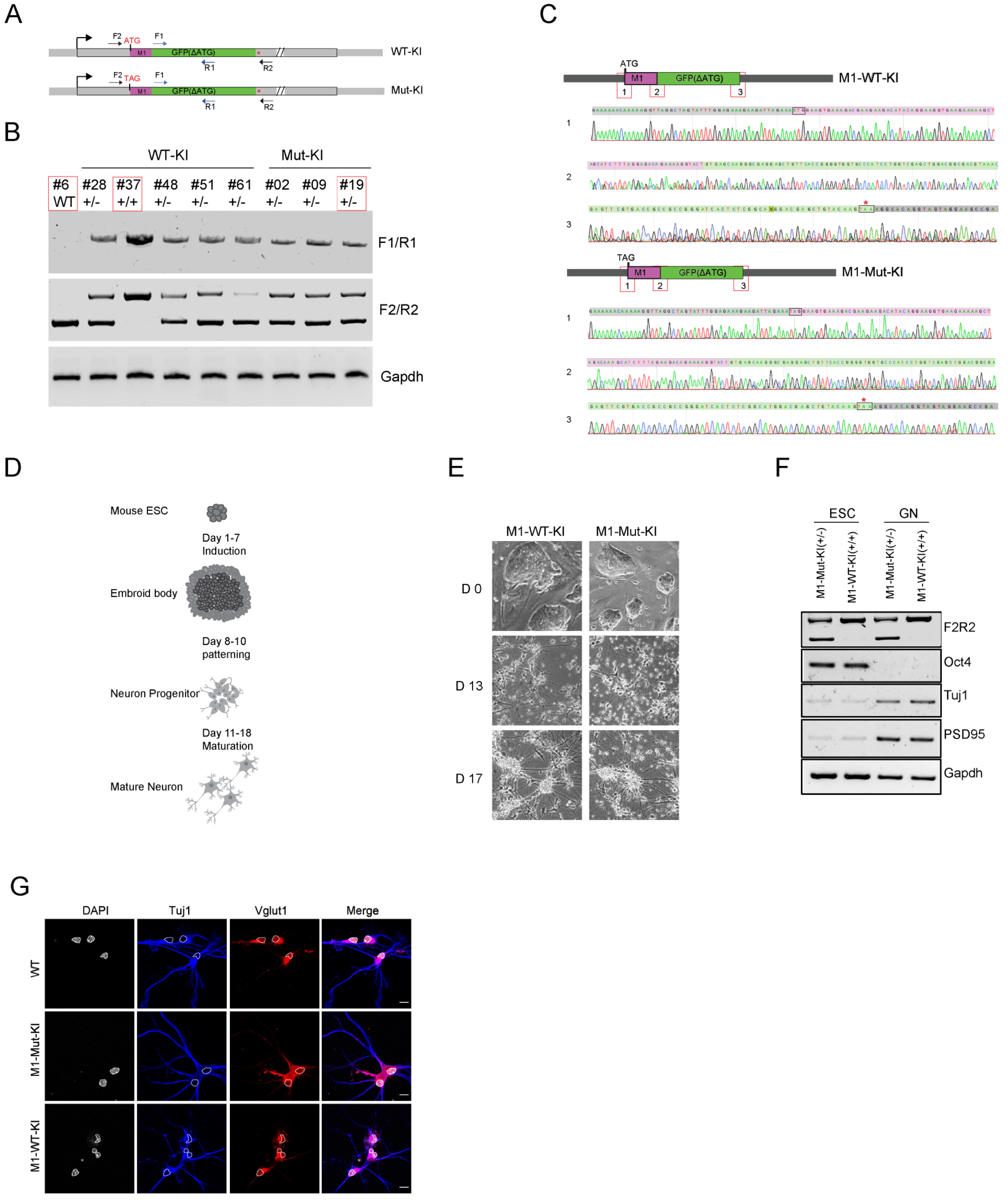

(A) Schematic of the Malat1 locus with GFP fused to the M1-ORF. (B) Genotype analysis using internal GFP targeting primers and the flanking primers targeting Malat1 region. The red boxes highlight clones selected for downstream differentiation experiments. One line homozygous for M1-WT-KI (#37) and one line heterozygous for M1-Mut-KI (#19) were chosen for further analysis. (C) Sanger sequencing of the Malat1 loci in the M1-WT-KI ES line (#37) and the M1-Mut-KI ES line (#19). Boxes 1, 2 and 3 highlight the start codon region of M1, the start codon region of GFP and stop codon region of GFP, respectively. (D) Diagram of the protocol for differentiating ES cells to glutamatergic neurons (GN). (E) Morphology of the M1-WT-KI and M1-Mut-KI ESC lines over differentiation. (F) RT-PCR analysis for pluripotency marker (Oct4) and neuronal markers (Tuj1 and PSD95) in the M1-WT-KI and M1-Mut-KI ESC lines and differentiated glutamatergic neurons. (G) Immuno-fluorescence assays for Tuj1 and vglut1 in glutamatergic neurons derived from the knock-in ES cells. Nuclei were highlighted with dash lines. Scale bar, 5 um.

**Supplemental Fig.8**: Reactivity and specificity of M1 antibody in immunoblot, immunoprecipitation and immunofluorescence assays.

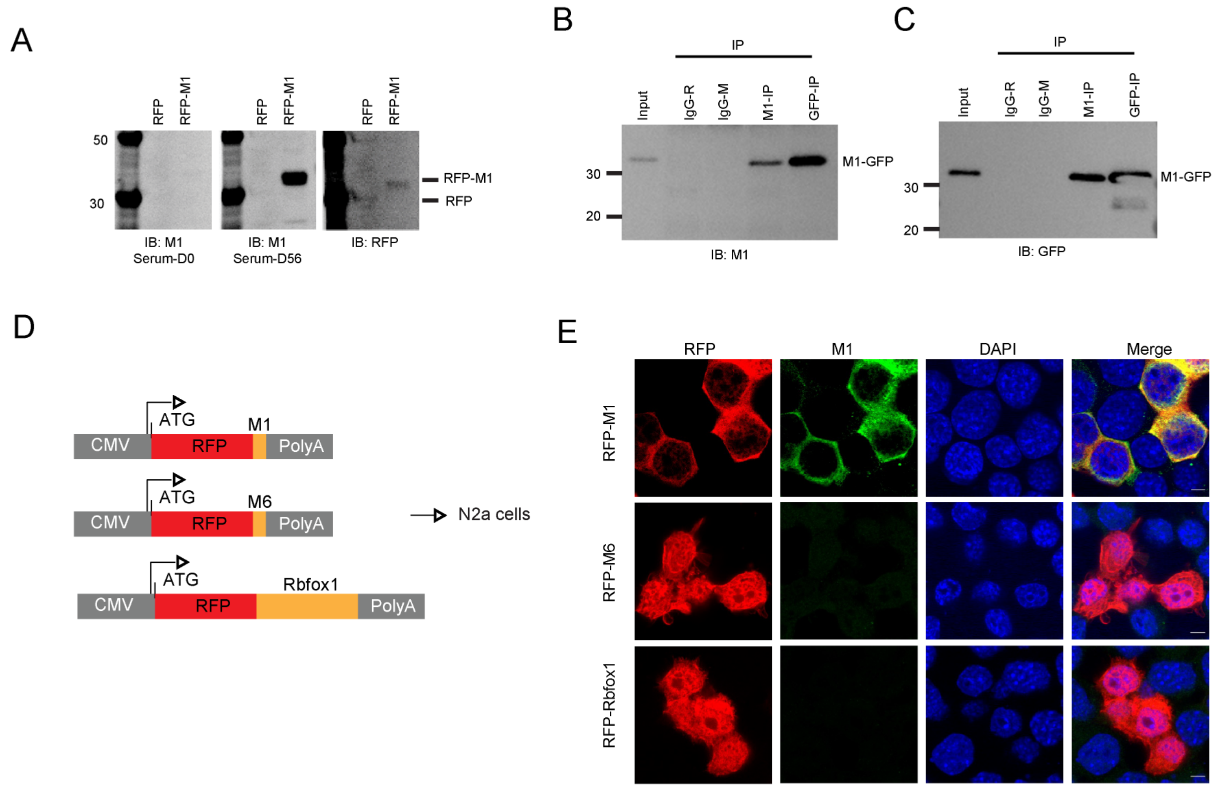


(A) M1 sera from day 0 and day 56 after immunization were applied to WB analysis in RPF and RFP-M1 expressing N2a cells. (B-C) Immunoprecipitation of M1-GFP with anti-M1 and anti-GFP antibodies in GFP-M1 expressing N2a cells. Rabbit and mouse IgG were negative controls. In (B), the input and IP samples were immunoblotted with M1 antibody. In (C), GFP antibody was used for immunoblot analysis. (D) Diagram of transient expression constructs with RFP fused to the M1 and M6 peptides, or to the Rbfox1 protein. (E) Immunofluorescence of transiently expressed RFP-M1, RFP-M6 and RFP-Rbfox1 protein in N2a cells. M1 antibody was used to detect RFP-M1 or endogenous M1. RFP was visualized by its own protein fluorescence.

### References

Gueroussov S, Gonatopoulos-Pournatzis T, Irimia M, Raj B, Lin Z-Y, Gingras A-C, Blencowe BJ. 2015. An alternative splicing event amplifies evolutionary differences between vertebrates. *Science* **349**: 868–873.

Saini H, Bicknell AA, Eddy SR, Moore MJ. 2019. Free circular introns with an unusual branchpoint in neuronal projections. *eLife* **8**: e47809.

Xiao W, Yeom K-H, Lin C-H, Black DL. 2023. Improved enzymatic labeling of fluorescent in situ hybridization probes applied to the visualization of retained introns in cells. *RNA* **29**: 1274–1287.
